# Immune Aging is an Independent Risk Factor for Cardiovascular Disease

**DOI:** 10.64898/2026.05.18.725733

**Authors:** Erik Feldman, Everton Jose Santana, Bettia Celestin, Nadav Golden, Shadi Bagherzadeh, Sofia Maysel, Kavita Mathi, Sarah Short, Megan Carroll, Shanon S. Sullivan, Martin Lukačišin, Xuhuai Ji, Yuval Klein, Oren Caspi, Patricia Kim Phuong Nguyen, William F. Fearon, Brian Kim, Svati Shah, Kenneth W. Mahaffey, Holden T. Maecker, Mark M. Davis, Neta Milman, Timothy J. Few-Cooper, Francois Haddad, Shai S. Shen-Orr

**Author notes:** Department of Genetics, Faculty of Natural Sciences, Comenius University, Ilkovičova 6, 84104, Bratislava, Slovakia (Martin Lukačišin). These authors jointly supervised this work (Neta Milman, Timothy J. Few-Cooper, Francois Haddad, Shai S. Shen-Orr).

## Abstract

Cardiovascular disease remains the leading cause of mortality, yet current clinical predictors miss substantial disease-risk. While the immune system contributes to this residual risk, its complexity has hindered broadly applicable, clinically scalable metrics of immune-state. Here, we establish the prognostic relevance of IMM-AGE, a system-level metric of immune-aging, to cardiovascular disease. We learn reference-free, low-dimensional representations of IMM-AGE across cell, protein, and mRNA measurements, enabling high-fidelity quantification across modalities, blood fractions, and platforms, including standard hospital flow cytometers. Among UK-Biobank participants, 56.9% of IMM-AGE variation remained unexplained by routine clinical measures, and across diverse cohorts totaling ∼48,000 individuals, elevated IMM-AGE was independently associated with future cardiovascular risk, intervention outcomes, and mortality. Moreover, incorporation of IMM-AGE into the PREVENT 10-year risk equation significantly improved risk stratification. These findings establish immune-aging as an independent biological dimension of cardiovascular disease-risk and support IMM-AGE as a practical tool for precision risk assessment.

## Introduction

Cardiovascular (CV) disease remains the leading cause of mortality worldwide, yet a substantial fraction of risk, often referred to as residual risk, remains unexplained^1,2^. Current clinical risk models, including the Framingham Risk Score, Pooled Cohort Equations, and more recently the PREVENT equations, estimate risk based on established factors such as age, sex, blood pressure, lipid levels, smoking status, and diabetes^3–5^. These tools are derived from large longitudinal cohorts and guide major clinical decisions, including initiation of statin therapy and antihypertensive treatment^6^. However, their predictive capacity is inherently limited to the variables they include. The implication is that biological processes not captured by these models remain effectively invisible, constraining our ability to fully account the risk of an individual to cardiovascular outcomes.

Cardiovascular disease is not solely a disorder of lipid accumulation or hemodynamic stress, but a process also shaped by sustained immune activation within vascular and cardiac tissues. Foundational work has established that immune pathways are integral to every stage of cardiovascular pathology, from the initiation of vascular injury to the progression of chronic disease^7,8^, including atherosclerosis^9,10^, myocardial remodeling^11^, and vascular dysfunction. Importantly, inter-individual variation in immune system state is substantial and structured^12,13^ raising the possibility that this heterogeneity could be harnessed as a distinct and clinically actionable risk factor for CVD. Indeed, several metrics of immune aging and health derived from cellular and molecular profiling have linked variation in immune state to aging, multimorbidity, all-cause mortality, and cardiovascular phenotypes^14–16^. However, these metrics have not been extensively evaluated against hard cardiovascular outcomes or integrated into established clinical risk-prediction frameworks and while high-dimensional proteomic studies have identified biomarkers that augment established cardiovascular risk equations such as PREVENT^17,18^, these approaches were not designed to quantify immune system state. Thus, despite growing evidence that immune state carries clinically relevant information, it remains largely absent from cardiovascular risk assessment. Closing this gap requires a measure of immune state that is not only biologically informative, but quantitative, reproducible, broadly applicable, clinically scalable, and integratable with the risk tools clinicians already use.

We previously developed IMM-AGE, the first high-resolution, system-level and quantitative metric of immune aging, derived from longitudinal remodeling of immune-cell composition^19^. IMM-AGE models this process as a continuous trajectory defined by coordinated shifts across 18 immune-cell populations, as measured by mass cytometry (CyTOF). Unlike clocks trained against chronological age or specific clinical endpoints, IMM-AGE reflects the developmental process of immune aging itself, capturing system-level changes in immune organization rather than a predefined disease phenotype. Moreover, IMM-AGE has been previously associated with cardiovascular phenotypes in FHS^19^, is based on quantitative features of circulating immune cells accessible from peripheral blood, and has recently been highlighted as the leading immune-aging measure for clinical-trial applications^20^. Together, these properties make IMM-AGE a particularly promising candidate for clinical translation, however, its original, high-dimensional CyTOF implementation limits scalability and accessibility for broader research and clinical use.

Here, we establish a translational framework for quantifying immune system state through IMM-AGE, and apply this to the study of cardiovascular disease. Specifically, we learn reference-free, low-dimensional representations of IMM-AGE across multiple molecular modalities including a reduced-marker, flow-cytometry-based implementation compatible with hospital flow cytometers with the potential for broad, clinical use. Using these representations, we systematically evaluated the relationship between immune aging and cardiovascular risk across population-based, disease-specific, and interventional settings. We first examined this relationship across 10 public cohorts comprising 885 individuals spanning chronic atherosclerotic, acute, valvular, pulmonary vascular and inflammatory cardiovascular disease subtypes, where IMM-AGE consistently tracked cardiovascular morbidity. In 43,608 UK Biobank (UKB) participants, conventional clinical risk factors explained less than half of the variation in IMM-AGE, with cardiovascular disease history accounting for only 1%, suggesting that immune aging is an independent and distinct dimension of biology rather than a restatement of known risk. Critically, this distinct, non-redundant signal was prognostically relevant: higher IMM-AGE independently predicted incident cardiovascular events over a median 13.5 years of follow-up and, when incorporated into the PREVENT framework, improved model fit and meaningfully refined individual risk classification beyond conventional risk factors and CRP. Finally, using the clinically oriented flow-cytometry implementation, IMM-AGE identified patients at risk of maladaptive recovery following transcatheter aortic valve replacement and distinguished future cardiovascular cases from closely matched controls in the prospective Baseline Health Study. Together, our findings position immune aging as a measurable and clinically tractable dimension of cardiovascular risk, and provide a path toward incorporating immune-system state into precision cardiovascular risk assessment.

## Results

### A translational molecular framework extends IMM-AGE across diverse cardiovascular disease settings

To assess the prognostic relevance of IMM-AGE in cardiovascular disease and facilitate its wider clinical translation, we developed a scalable, translational framework to extend IMM-AGE beyond its original measurement setting. The SELA cohort is a unique resource comprising high-resolution, densely sampled longitudinal data of healthy adults, within which we identified the patterns of immune cell compositional shift over age and developed the IMM-AGE metric. While the size of the SELA cohort precludes direct inference of clinical consequences, its multi-omic richness enables the identification of predictive features in other modalities that serve as proxy representations of IMM-AGE, enabling its evaluation across clinical conditions in public and proprietary datasets (**Fig. 1A**).

**Figure 1.**
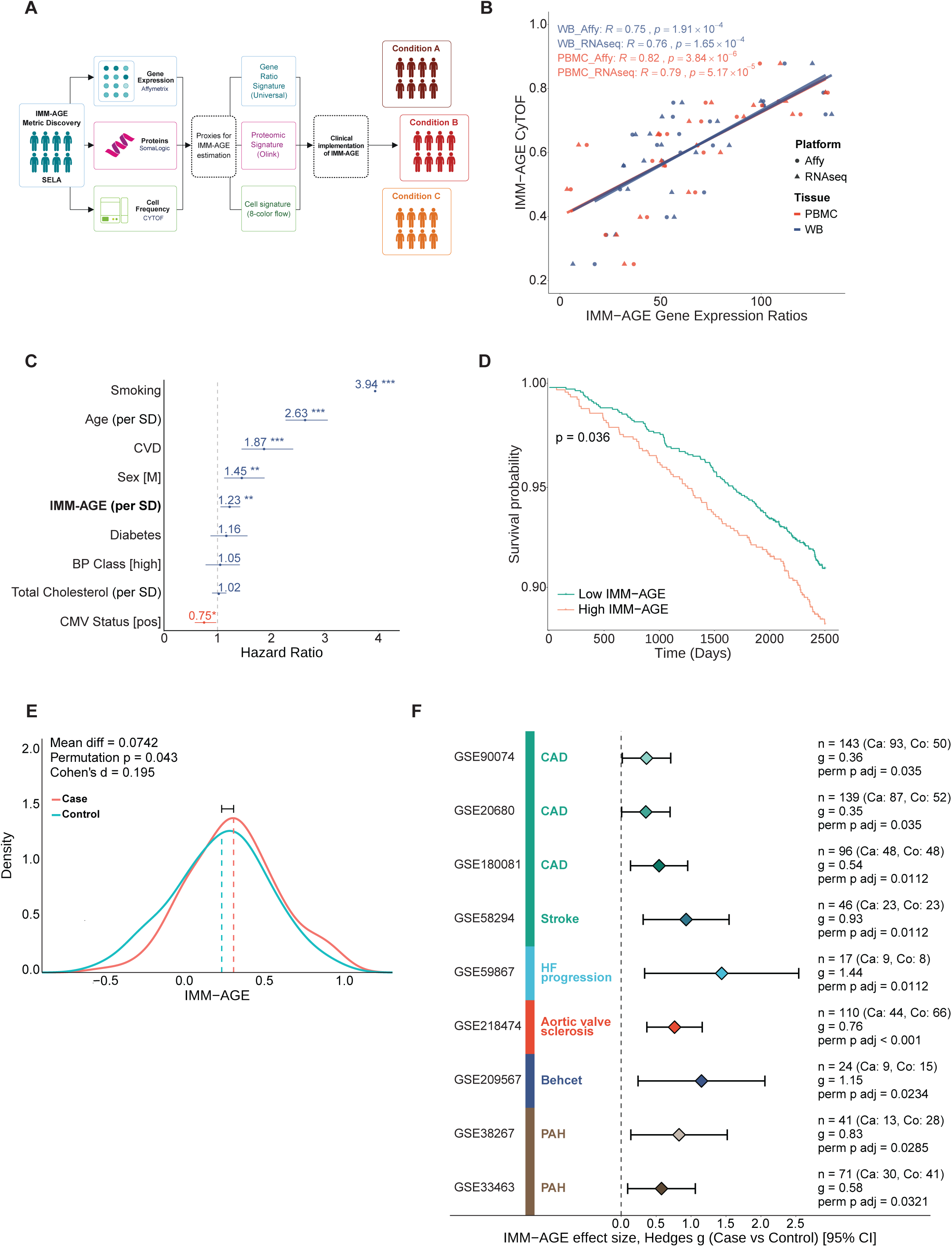
IMM-AGE is a quantifiable metric of cardiovascular morbidity and mortality risk. **(A)** Schematic of a molecular-clinical, translational framework designed to translate data from a high-resolution, longitudinal multi-omic study of immune aging into scalable molecular proxies and validated fine-grained clinical associations in large cross-sectional cohorts. **(B)** Ratio-based IMM-AGE is preserved across blood fractions and transcriptomic platforms. Scatter plot comparing IMM-AGE estimated from gene-ratios to CyTOF-derived IMM-AGE scores in whole blood (WB) and PBMC samples, profiled by either Affymetrix microarray or RNA-seq. Each point represents a sample; colors indicate tissue source and point shapes indicate transcriptomic platform. Linear regression models were fitted separately by tissue and are shown as solid lines. Correlation coefficients across tissue– platform combinations ranged from R=0.753 to 0.824 (all *P<0.001*).**(C)** Independent prognostic value of IMM-AGE in the Framingham Heart Study. Forest plot of hazard ratios from a multivariable Cox model including 2,131 participants and 297 mortality events. IMM-AGE remained independently associated with mortality following adjustment for conventional cardiovascular risk factors (HR = 1.23, p = 7.7×10^−3^). Abbreviations: CVD: History of cardiovascular disease; BP Class [High]: sysBP≥130 OR diaBP≥85. **(D)** Kaplan–Meier curves showing overall survival in the Framingham cohort, as stratified by IMM-AGE status. Participants with high IMM-AGE (above the 60th percentile; red) had lower overall survival than those with low IMM-AGE (green) (*N* = 2,131; log-rank *P* = 0.036). **(E)** Distribution of IMM-AGE scores in patients with coronary artery disease and age- and sex-matched controls. Patients with coronary artery disease had higher IMM-AGE scores than matched controls (mean difference = 0.0786; paired permutation test, *P* = 0.0434). Dashed lines indicate group means. Data are from GSE20681. **(F)** Forest plot summarizing differences in IMM-AGE across ten independent transcriptomic datasets spanning cardiovascular and cardiometabolic disease. Effect estimates (Hedges’ *g*) and 95% confidence intervals are shown for each dataset. Higher IMM-AGE scores were observed in cases than controls across all cohorts including those detailing coronary artery disease, cerebrovascular events (stroke), post-myocardial infarction heart-failure progression, aortic valve sclerosis, vascular Behçet’s syndrome and pulmonary arterial hypertension. GEO accession numbers and sample sizes are shown.

We first focused on a transcriptomic proxy, where the breadth of available datasets provided an opportunity to test both its portability across diverse cohorts and assay conditions but also obtain an initial assessment of the relationship between IMM-AGE and cardiovascular phenotypes. This breadth, however, comes with substantial heterogeneity in experimental design. In particular, transcriptomic datasets reporting cardiovascular outcomes span different blood fractions and profiling platforms. A proxy is only useful if its readout is determined by the underlying biology rather than by the assay. We therefore learned a signature across multiple assay conditions and constructed a transcriptomic IMM-AGE signature in SELA from pairwise ratios between genes associated with the immune populations underlying IMM-AGE. By capturing the relative abundance of two transcripts, these ratios are self-normalizing and largely invariant to scaling and batch effects across blood fractions and platforms. Specifically, in 20 SELA participants held out from signature training and independently profiled by RNA-seq and Affymetrix microarrays across whole-blood and PBMC samples, gene ratio-based IMM-AGE closely tracked CyTOF-derived estimates with high fidelity across all fraction and platform combinations (**Fig. 1B**, Spearman ⍴>0.753 and p<1.9×10⁻^4^ across all comparisons; **Supplementary Fig. 1A-C**, **Table 1**; see Methods). Moreover, in the Framingham Heart Study (FHS), where the original 56-gene IMM-AGE signature demonstrated clinical relevance^19^, ratio-based scores strongly resembled the original estimates (n = 2,418; (**Supplementary Fig. 1D;** Spearman ⍴=0.72, p<2.2×10⁻¹⁶), and, in 2131 individuals with complete data, remained independently associated with all-cause mortality in a fully adjusted Cox model (297 deaths; median follow-up 8.2 years; **Fig. 1C**; HR = 1.23, p=7.7×10^−3^ per SD; see Methods) while also stratifying overall survival (**Fig. 1D**; log-rank p=3.6×10^−2^; see Methods). Together, these findings support gene ratio-based IMM-AGE as a portable transcriptomic proxy that preserves the clinical signal of the original metric.

Cardiovascular disease spans chronic, acute, valvular, inflammatory and metabolically driven states arising in distinct biological contexts^21–23^, and as such we asked whether the IMM-AGE signal seen in FHS extends across this landscape. To address this, we analyzed 10 independent gene-expression cohorts (N=885 individuals), drawn from the CVD atlas^24^, and Gene Expression Omnibus, for which phenotypic annotation, defined clinical outcomes and sufficient sample sizes were available (see Methods). Across all cohorts, higher IMM-AGE consistently marked disease or adverse cardiovascular phenotypes (**Fig. 1E,F; Supplementary Table 2**, see Methods). In chronic atherosclerosis, individuals with significant coronary stenosis had higher IMM-AGE than age- and sex-matched controls (**Fig. 1E**, perm p = 4.3×10⁻², **Supplementary Table 2**, GSE20681; see Methods), with similar signal observed in three additional, non-matched pre-catherization cohorts (**Fig. 1F**; adj p 3.5×10⁻² for both GSE20680 and GSE90074, and 1.1×10⁻² for GSE180081, **Supplementary Table 2**; see Methods). In acute, ischemic disease, baseline IMM-AGE distinguished patients presenting with myocardial infarction who subsequently progressed to heart failure (HF) from those who remained free of HF (**Fig. 1F**; adj p=1.1×10⁻², GSE59867; **Supplementary Table 2;** see Methods), and was elevated in patients experiencing cardioembolic stroke prior to treatment (**Fig. 1F**; adj p=1.1×10⁻², GSE58294). Importantly, this pattern was not restricted to classical atherosclerotic or ischemic presentations. IMM-AGE was elevated in patients with aortic valve sclerosis accompanying acute myocardial infarction **(Fig. 1F**; adj p < 1×10⁻³, GSE218474; **Supplementary Table 2;** see Methods), and in Behçet’s syndrome - a clinically relevant model of inflammation-driven cardiovascular disease - where IMM-AGE elevation was confined to the vascular subtype (Fig. 1F; adj p = 2.3×10⁻², GSE209567) that confers greater cardiovascular risk but not in the mucocutaneous, ocular or complex disease variants (g = 0.16, p = 0.631). Finally, IMM-AGE was elevated in pulmonary arterial hypertension across two independent cohorts (Fig. 1F; adj p = 2.9×10⁻² and 3.2×10⁻²; GSE38267 and GSE33463). Together, these findings indicate that the association between IMM-AGE and cardiovascular morbidity extends across a broad spectrum of cardiovascular disease rather than being confined to any single manifestation.

### IMM-AGE captures residual cardiovascular risk in the UK Biobank

While publicly available gene-expression datasets revealed a broad association between IMM-AGE and cardiovascular disease, they provided little insight into how immune-aging, as quantified by IMM-AGE, relates to conventional clinical and cardiovascular risk factors owing to their limited phenotypic and demographic data. To address this gap in a population setting, we turned to UK Biobank (UKB) participants with both multiplex serum proteomics and deep clinical and phenotypic annotation^25^. Subsequently, we applied the SELA translational framework to develop a serum proteomic proxy of IMM-AGE compatible with Olink data, enabling its estimation in the UKB (**Supplementary Table 3**, see Methods).

Using this surrogate, we analyzed 43,608 UKB participants with proteomic data and decomposed variation in IMM-AGE across demographic features, inflammatory and infectious burden, metabolic and lifestyle exposures, kidney function, and medical history of cardiovascular and non-cardiovascular disease (**Fig. 2A**; see Methods). Together, these variables explained 43.1% of the total variation in IMM-AGE scores. As expected, Age was the largest single contributor (10.7%), followed by BMI, CRP and eGFR (10.4%, 7.8% and 5.1%, respectively). Notably, a history of CVD accounted for only 1% of the explained variance, suggesting that the association between IMM-AGE and cardiovascular morbidity is not simply driven by pre-existing disease. Accordingly, 56.9% of variation in IMM-AGE remained unexplained by standard clinical descriptors, suggesting that IMM-AGE partially reflects conventional clinical features but retains substantial, additional information beyond that normally recorded.

**Figure 2.**
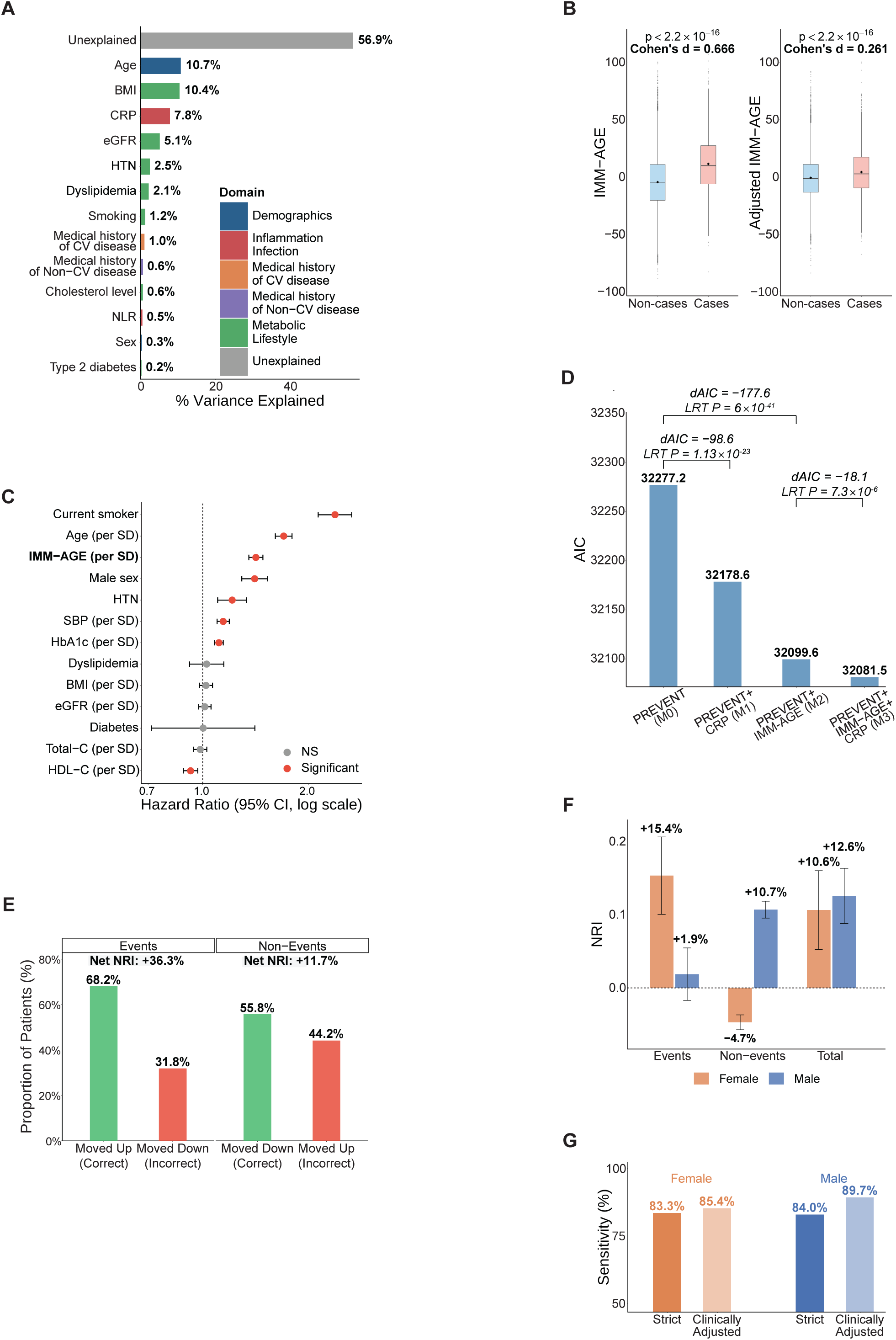
IMM-AGE captures residual cardiovascular risk in the UK Biobank cohort. (A) **(A)** Variance decomposition of IMM-AGE across clinical domains in 43,608 UK Biobank participants. Bars show the proportion of IMM-AGE variance explained by individual demographic, inflammatory, metabolic, lifestyle, and disease-related factors. Collectively, measured clinical factors explained 43.1% of total variance (R^2^=0.431), while 56.9% remained unexplained. HTN, hypertension defined by blood-pressure treatment. **(B)** IMM-AGE is elevated in individuals who subsequently develop cardiovascular events. Baseline IMM-AGE (left) and multivariable-adjusted IMM-AGE (right) are shown for participants who were event-free at baseline but developed a cardiovascular event during follow-up (“cases”) versus those who remained event-free (“non-cases”). IMM-AGE remained significantly elevated in cases both before and after adjustment for conventional clinical risk factors (two-sided *t*-test, *P* < 2.2 × 10^−16^; Cohen’s *d* = 0.666 and 0.261, respectively). **(C)** IMM-AGE is an independent risk factor for cardiovascular morbidity. **(C)** IMM-AGE is an independent risk factor for cardiovascular morbidity. Multivariable Cox proportional hazards model showing that higher baseline IMM-AGE was independently associated with incident cardiovascular events (HR = 1.417 per SD; 95% CI, 1.353–1.484; *P* = 1.36 × 10^−49^), alongside established cardiovascular risk factors. eGFR was estimated using the CKD-EPI 2021 equation; HTN and dyslipidemia were defined by blood-pressure and lipid-lowering treatment, respectively. **(D)** IMM-AGE improves model fit beyond PREVENT, outperforming CRP. Akaike information criterion (AIC) values are shown for four nested Cox models using the PREVENT logit as a fixed offset. Brackets indicate differences in AIC (ΔAIC) between models and the corresponding likelihood-ratio test (LRT) *P* values. M0, PREVENT; M1, PREVENT + CRP; M2, PREVENT + IMM-AGE; M3, PREVENT + IMM-AGE + CRP. **(E)** Continuous NRI after incorporating IMM-AGE into PREVENT. Addition of IMM-AGE shifted predicted risk upward more often than downward among participants who experienced events, and downward more often than upward among those who remained event-free, yielding event and non-event NRI values of +36.3% and +11.7%, respectively. Events, *n* = 1,680; non-events, *n* = 28,454. **(F)** IMM-AGE improves PREVENT-based categorical risk reclassification. Categorical net reclassification improvement (NRI) after addition of IMM-AGE to PREVENT is shown separately for women and men. IMM-AGE yielded significant total NRI in both sexes, driven predominantly by upward reclassification among participants who subsequently experienced cardiovascular events. **(G)** Clinically adjusted reclassification preserves treatment eligibility while improving sensitivity. Sensitivity of PREVENT-based risk reclassification after addition of IMM-AGE is shown under strict and clinically adjusted definitions. Under the clinically adjusted definition, downward reclassification was not considered adverse when lipid-lowering treatment eligibility was preserved.

Given the extent of this orthogonality, we reasoned IMM-AGE may further discern individuals at elevated cardiovascular risk before overt disease develops. We therefore tested whether IMM-AGE at study enrollment (hereafter, “UKB baseline”) predicted incident cardiovascular events among participants initially free of cardiovascular disease (**Supplementary Tables 4-5**). We observed that individuals who subsequently developed atherosclerotic cardiovascular disease (ASCVD) or heart failure (HF) (n=2760) had higher IMM-AGE values at UKB baseline than those who remained event-free during follow-up (**Fig. 2B**; t-test, p<2.2×10⁻¹⁶). This difference remained significant following adjustment for age, HbA1c, BMI, smoking status, systolic blood pressure, antihypertensive treatment, lipid-lowering treatment, type-2 diabetes, eGFR, HDL, and total cholesterol (**Fig. 2B**; t test, p<2.2×10⁻¹⁶). In time-to-event analyses, higher IMM-AGE at UKB baseline was independently associated with incident cardiovascular events in a multivariable Cox proportional hazards model (**Fig. 2C**; HR = 1.417 per SD; 95% CI 1.353-1.484; P = 1.36×10^−49^; see Methods). IMM-AGE is, therefore, able to predict incident cardiovascular events independently of pre-existing clinical risk factors.

Cardiovascular risk is now routinely estimated using multivariable models that integrate established clinical risk factors. We therefore asked whether incorporating IMM-AGE into these risk models could improve cardiovascular risk stratification. We focused on PREVENT, the current model of the American Heart Association (AHA), which integrates demographic, metabolic, renal, blood-pressure, lipid and diabetes variables to estimate 10-year cardiovascular risk and guide preventive treatment, and benchmarked the incremental contribution of IMM-AGE against CRP - an established marker of inflammatory burden - using UKB participants as before. Adding IMM-AGE to PREVENT yielded a greater improvement in model fit than by adding CRP, as reflected by a larger reduction in the Akaike information criterion (AIC) (**Fig. 2D**; ΔAIC = −177.6 vs. −98.6; **Supplementary Tables 6-7**). Moreover, IMM-AGE remained informative after adjustment for CRP, whereas adding CRP to PREVENT + IMM-AGE produced only a modest additional improvement in model fit (ΔAIC = −18.11; P = 7.3 × 10⁻⁶). We further observed that IMM-AGE produced a modest gain in global rank concordance (**Supplementary Fig. 2A** and **Supplementary Table 8**) but significantly increased time-dependent discrimination at the clinically relevant 10-year horizon **(Supplementary Fig. 2B**; AUC 0.726 to 0.741; ΔAUC = +0.015; P = 1.4 × 10⁻¹¹, **Supplementary Table 9**). Continuous net reclassification improvement (NRI) analysis, which assesses whether predicted risk shifts in the appropriate direction for individuals with and without events, showed that IMM-AGE increased predicted risk in 68.2% of participants who developed cardiovascular events and decreased it in 55.8% of event-free participants, yielding event and non-event NRI values of +36.3% and +11.7%, respectively (**Fig. 2E**; total NRI = +48%). Thus, IMM-AGE refines cardiovascular risk estimation beyond PREVENT and CRP.

As PREVENT-based risk estimates inform treatment decisions (e.g. statin prescription) using clinically defined, sex-specific categories - low (<3%), borderline (3-5%), intermediate (5-10%), and high (≥10%) - we next asked whether adding IMM-AGE meaningfully re-classified individuals rather than merely shifting predicted risk percentages alone. Indeed, IMM-AGE produced significant categorical net reclassification improvement in both women and men, with total NRI values of 10.6% and 12.6%, respectively (**Fig. 2F**; **Supplementary Fig. 2C**; both P < 0.001). Among participants who subsequently developed cardiovascular events, IMM-AGE preferentially shifted predicted risk upward, yielding event NRI values of 15.4% in women and 1.9% in men, a significant sex-specific difference in events and non-events (Δ=13.6 percentage points, 95% CI 13.2-14.2, and Δ=-15.3 percentage points, 95% CI −18.0 to −12.6 in events and non-events respectively, all P<0.005 by bootstrap difference and permutation testing of sex-blind random splits; **Supplementary Fig. 2D**). Importantly, these upward shifts were concentrated in clinically actionable intermediate-and high-risk PREVENT strata (**Supplementary Fig. 2E**). Overall, event cases showed median per-person risk increases of 2.5 and 3.3 percentage points in women and men, respectively (**Supplementary Fig. 2F**). Among non-events, IMM-AGE primarily shifted low- and borderline-risk individuals downward, although a subset of high-risk non-events moved upward, consistent with elevated immune-risk profiles despite no observed events (**Supplementary Fig. 2E-F**). Using treatment-aligned definitions, IMM-AGE preserved high sensitivity and specificity across sexes (**Supplementary Fig. 2G**). In a clinically adjusted analysis, where downward shifts remaining within the intermediate-risk category were not penalized as these individuals would retain the same inferred eligibility for lipid-lowering therapy, sensitivity increased to 85.4% in women and 89.7% in men (**Fig. 2G**). Taken together, these findings show that IMM-AGE adds prognostic information to PREVENT beyond CRP - improving model fit and refining individual-level risk both continuously and across clinically relevant risk categories - while capturing dimensions of cardiovascular risk not represented by conventional clinical factors and further implicating immune aging in cardiovascular disease.

### Clinical translation of IMM-AGE is feasible with standard flow cytometry

While our transcriptomic and proteomic proxies provide powerful tools to evaluate IMM-AGE at scale and establish its relevance to CVD across diverse public cohorts and UKB, neither offers a straightforward path to routine clinical implementation owing to cost, complexity and limited accessibility. Because IMM-AGE is fundamentally rooted in immune-cell composition, we reasoned that standard clinical flow cytometry could provide a practical alternative, combining scalability, lower cost and rapid turnaround. The key challenge, however, is dimensionality: CyTOF enables the simultaneous measurement of more than 40 parameters in a single sample, whereas standard clinical flow cytometry is considerably more constrained. Nevertheless, because IMM-AGE reflects coordinated shifts across immune-cell populations rather than the abundance of any single population, we hypothesized that its underlying signal could be preserved in a low-dimensional assay if the key relationships among immune subsets were retained. We therefore asked whether IMM-AGE could be recovered using a substantially reduced set of cell-surface markers.

To identify such a reduced panel, we used SELA CyTOF data to exhaustively evaluate all eight-marker subsets of the original CyTOF panel. For each candidate panel, we re-estimated IMM-AGE scores for all individuals in the original SELA study^19^ (**Fig. 3A**; **Supplementary Fig. 3A**; see Methods). We subsequently ranked panels by their ability to preserve sample ordering along the full-panel IMM-AGE trajectory and, after translation into gene-expression signatures, to retain similar prognostic performance in FHS to that observed in the original IMM-AGE study (**Fig. 3A**; see Methods). This screen identified three optimized panels that closely reproduced full-panel, CyTOF-based IMM-AGE while maintaining prognostic performance (**Fig. 3B**; Kendall’s τ > 0.95 for reproducibility; C-index > 0.719 for prognostic performance; **Supplementary Table 10**; see Methods).

**Figure 3.**
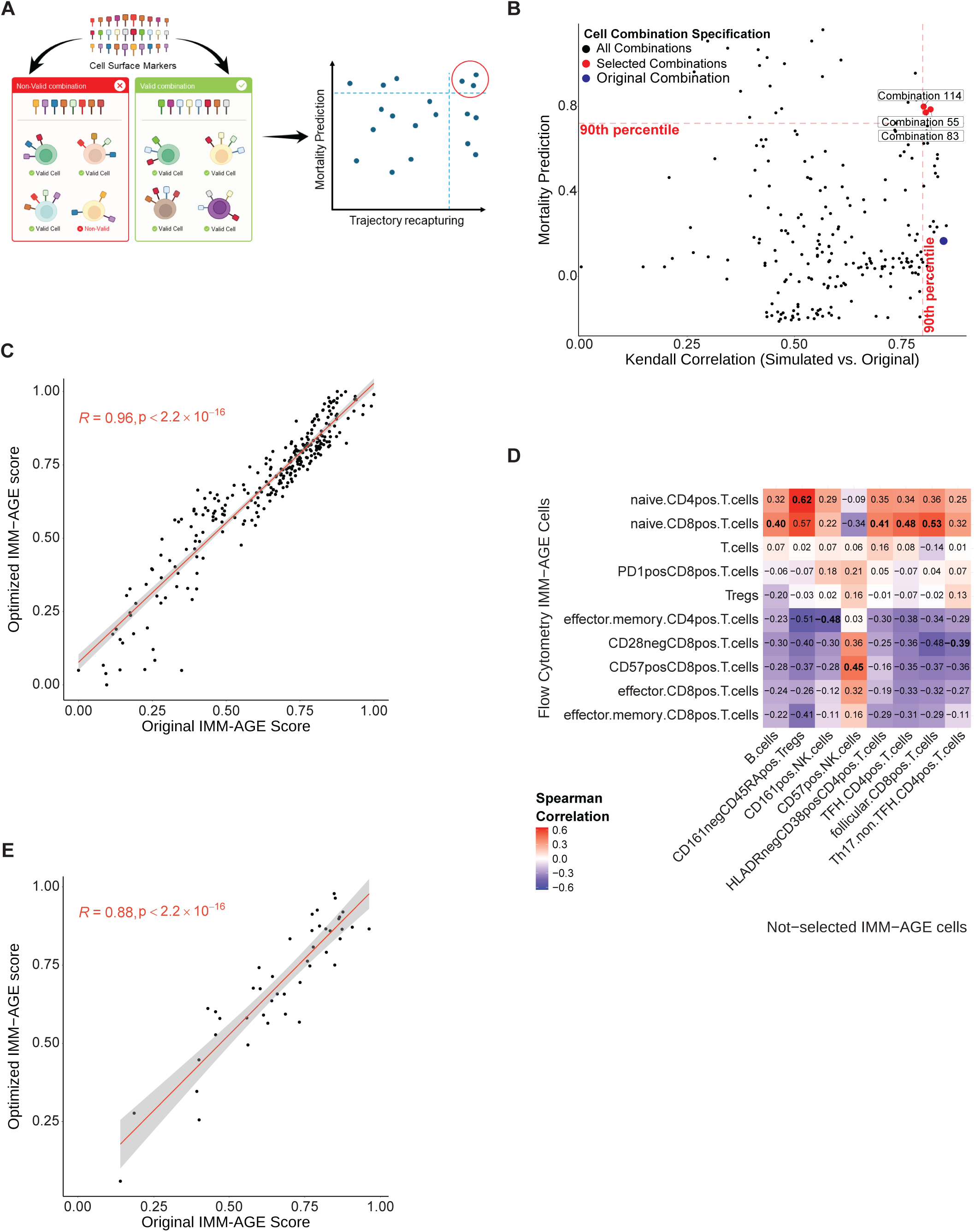
A scalable framework for monitoring immune aging in standard clinical settings. **(A)** Identification of optimized marker panels for clinical translation. Schematic illustrating the computational framework used to reduce the original, high-dimensional CyTOF-based IMM-AGE to lower-dimensional panels suitable for conventional flow cytometry. Optimized configurations were selected to preserve ordering along the original IMM-AGE trajectory and retain predictive value for mortality in the Framingham Heart Study. **(B)** Selection of high-fidelity marker panels based on trajectory preservation and mortality prediction. Scatter plot showing all simulated eight-marker panels ranked by their ability to recapitulate the original IMM-AGE trajectory (Kendall’s τ, x-axis) and their prognostic performance for mortality in the Framingham Heart Study (C-index, y-axis). Selected eight-marker panels are shown in red, and the original CyTOF-based configuration in blue. Dashed lines indicate the selection thresholds for each metric. **(C)** Estimated reduced-panel IMM-AGE recapitulates the full CyTOF signal in the discovery cohort. Scatter plot comparing IMM-AGE scores derived from the original full CyTOF panel (x-axis) with scores estimated *in silico* using the selected optimized eight-marker panel (y-axis; Combination 55) in the SELA discovery cohort. Reduced-panel estimates showed strong concordance with the original IMM-AGE scores (Pearson R=0.96, *P* < 2.2 × 10^−16^). **(D)** Selected flow-based IMM-AGE populations capture information about broader immune-cell changes. Heatmap showing Spearman correlations between cell populations included in the reduced flow cytometry IMM-AGE panel and non-selected IMM-AGE-related populations. **(E)** Experimental validation of the reduced IMM-AGE panel. Scatter plot comparing IMM-AGE scores derived from the original mass cytometry (CyTOF) measurements (x-axis) with scores generated experimentally using the optimized eight-marker flow cytometry panel (y-axis) in a newly assayed subset of SELA participants (*n* = 46). Flow-derived IMM-AGE showed strong concordance with the original CyTOF-derived scores (Pearson R=0.88, *P* = 6.8 × 10^−15^).

Notably, all top ranked reduced IMM-AGE panels were comprised exclusively of T-cells subsets, despite the original IMM-AGE profile also spanning B-cell and myeloid lineages, and shared a common backbone of naïve CD4 T cells, naïve CD8 T cells, effector CD8 T cells, effector memory CD8 T cells and effector memory CD4 T cells, differing only in the inclusion of CD28^−^ CD8 T cells, CD57^+^ CD8 T cells, regulatory T cells and PD1^+^ CD8 T cells. For all subsequent analyses, we selected the panel containing CD28^−^ CD8 T cells in addition to the shared backbone. Reduced-panel IMM-AGE estimates closely tracked the original scores across the full IMM-AGE trajectory (Spearman’s ρ = 0.96), with minimal deviation from full-panel measurements (**Fig. 3C; Supplementary Fig. 3B**). Notably, the selected subsets captured a broader co-correlation structure spanning non-selected immune populations, supporting the idea that reduced-panel performance reflects preservation of higher-order immune organization rather than an arbitrary choice of markers (**Fig. 3D**). Consistent with this, and to assess signature flexibility, we systematically substituted each selected subset with alternative populations and quantified the resulting loss of fidelity, which we observed to be dependent upon the broader correlation structure rather than any single marker (**Supplementary Fig. 3C**). Next, we tested the selected panel in 46 additional SELA participants newly profiled by flow cytometry (**Supplementary Fig. 4A,B**; **Supplementary Table 11**; see Methods). Projection of these samples onto the original IMM-AGE trajectory yielded flow-based scores that closely agreed with the corresponding CyTOF-derived estimates (**Fig. 3E; Supplementary Fig. 4C**, Spearman ⍴=0.88, p<1.1×10⁻¹⁴; see Methods), consistent with the strong concordance between flow and CyTOF measurements of the selected immune populations (**Supplementary Fig. 4D**). Sequential exclusion of individual cell populations had only modest effects on IMM-AGE estimation, suggesting the estimate is robust (**Supplementary Fig. 4E**). Taken together, these results show that the higher-order immune state captured by IMM-AGE can be preserved in a low-dimensional assay, supporting its implementation by standard flow cytometry in routine clinical and hospital settings.

### IMM-AGE predicts maladaptive early cardiac recovery in an intervention setting

Having established a candidate, clinically scalable flow-cytometry implementation of IMM-AGE, we next sought to directly test its utility in real-world cardiovascular settings. Interventional settings provide a particularly compelling and informative test, asking whether pre-existing immune state can predict response to a defined clinical perturbation rather than simply associate with cardiovascular disease in the long-term.

We, therefore, first examined patients undergoing transcatheter aortic valve replacement (TAVR), where pre-procedural risk stratification currently relies on cardiac imaging, clinical status and routine laboratory variables, with little consideration given to the immune state of a patient. Notably, pre-existing clinical and demographic models provide only moderate discriminative power for key outcomes^26^, and while prior studies have linked pre-existing inflammatory phenotypes to adverse remodeling and mortality after TAVR^27^ none have sought to quantify or utilize this observation. We therefore tested whether flow-based IMM-AGE measured prior to TAVR could identify patients at risk of maladaptive recovery. Specifically, we analyzed 75 patients with symptomatic severe aortic stenosis who were at high surgical risk or deemed inoperable and underwent longitudinal assessment before TAVR (hereafter, “TAVR baseline”) at 1 month and at 1 year following the procedure, including blood sampling, echocardiography, and catheterization-based assessment (**Fig. 4A**; **Supplementary Table 12-13**; see Methods)^28^. To define maladaptation, we characterized cardiac state using four echocardiographic parameters capturing cardiac structure and function (LVEF, GLS-endo, LVMI and RVSP), and assessed these phenotypes at TAVR baseline and 1 month post-procedure^26,27^. Using these criteria, we classified 28 patients at TAVR baseline as having an unfavourable echocardiographic phenotype and 47 as favourable (**Supplementary Table 14**; **Supplementary Fig. 5A**). At 1 month, 23 were classified as showing maladaptive remodeling and 52 showed an adaptive response (**Supplementary Table 15**; **Supplementary Fig. 5B**), while by one-year follow-up, 11 patients had died (**Fig. 4B**). Notably, 1-month echocardiographic phenotypes were associated with subsequent mortality, with 30% of patients in the maladaptive group dying within 1 year, whereas 90% of those in the adaptive group survived (**Fig. 4C**; permutation test, p=4.4×10^−2^). By contrast, echocardiographic phenotypes at TAVR baseline were not associated with mortality (**Fig. 4C**; permutation test, P = 0.28). In contrast to this, patients who exhibited maladaptive ventricular remodeling at 1 month had significantly higher pre-procedural IMM-AGE scores at TAVR baseline than those with an adaptive response (**Fig. 4D**; t-test, p=6×10^−3^) (**Supplementary Fig. 5C-E**). This association remained significant in a multivariable logistic regression model adjusting for demographic, clinical and inflammatory covariates (**Fig. 4E**, log-odds=1.87, p=1.3×10^−2^). Notably, each standard deviation increase in IMM-AGE was associated with a 6.49-fold increase in the odds of a maladaptive 1-month echocardiographic phenotype, making it one of the strongest independent signals in the model alongside male sex (log-odds= 2.9, p=7×10^−3^), whereas most conventional variables, including BMI, metabolic syndrome and cardiovascular comorbidity, showed negligible or non-significant effects following adjustment.

**Figure 4.**
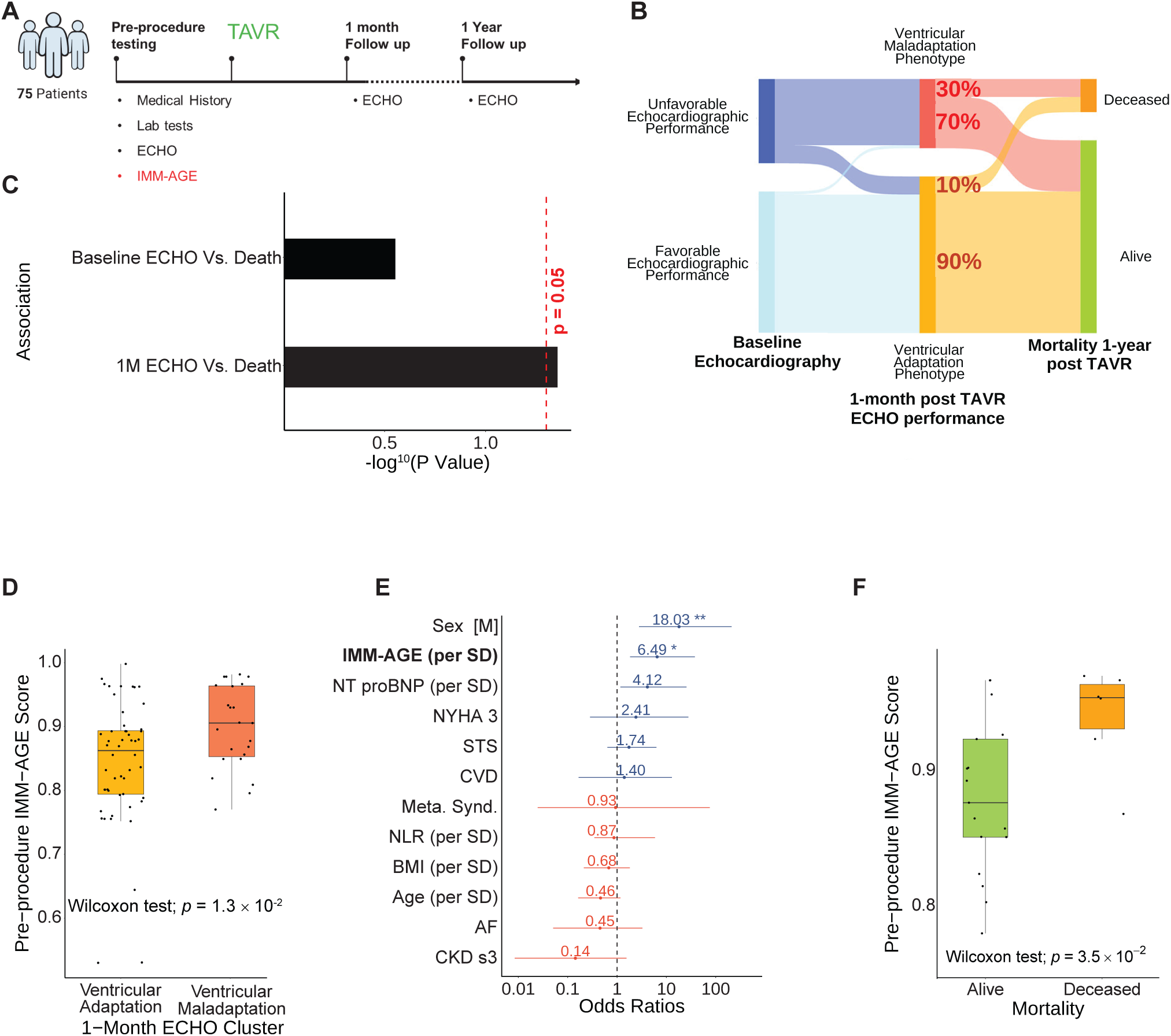
IMM-AGE is an independent predictor of post-TAVR ventricular adaptation and mortality. **(A)** Schematic of the prospective study design involving 75 patients undergoing transcatheter aortic valve replacement (TAVR). Assessments were performed at baseline (pre-procedure), 1 month, and 1 year after TAVR, and included clinical history, echocardiography, and IMM-AGE measurement using the optimized flow cytometry panel. **(B)** Echocardiographic trajectories from baseline through 1-month ventricular adaptation to 1-year survival. Sankey diagram illustrating the progression from baseline echocardiographic phenotypes through 1-month echocardiographic clusters to 1-year survival outcomes. **(C)** One-month ventricular adaptation, but not baseline echocardiographic phenotype, is associated with 1-year mortality. Bar plot showing the strength and significance of the association between echocardiographic phenotypes and 1-year mortality at baseline and 1 month after TAVR. Only the 1-month phenotype was significantly associated with 1-year mortality (permutation *P* = 0.044). **(D)** Baseline IMM-AGE is associated with subsequent 1-month ventricular adaptation. Boxplot comparing pre-procedural IMM-AGE scores between patients who subsequently developed adaptive versus maladaptive phenotypes at 1-month. Higher baseline IMM-AGE was associated with a maladaptive response (*t*-test, *P* = 0.006). **(E)** Baseline IMM-AGE independently predicts 1-month ventricular maladaptation. Odds ratio plot showing associations between baseline clinical factors and maladaptive ventricular response 1 month after TAVR. IMM-AGE was among the strongest independent predictors of maladaptation (OR = 6.49 per SD; *P* = 0.013) after adjustment for age, sex, NT-proBNP, and other clinical comorbidities. For continuous variables, ORs are reported per 1-SD increase. **(F)** Baseline IMM-AGE stratifies 1-year mortality among patients with maladaptive 1-month echocardiographic responses. Among patients classified as maladaptive at 1 month, those who died within 1 year had higher pre-procedural IMM-AGE scores than survivors (two-sided Wilcoxon test, *P* = 0.035).

To test whether this signal was simply a restatement of risk already captured by other, simpler means, we challenged it in three ways: (i) residualizing IMM-AGE against demographic, clinical and inflammatory factors left it associated with maladaptive recovery (**Supplementary Fig. 5F;** Wilcoxon test p=3.44×10^−2^, see Methods), indicating that elevated immune age precedes impaired recovery rather than merely reflecting these markers; (ii) IMM-AGE remained associated with maladaptive recovery after adjustment for baseline echocardiographic phenotypes, itself a strong determinant of recovery (**Supplementary Fig. 5G**; IMM-AGE log-odds = 1.322, p = 4.9×10⁻²; baseline phenotype log-odds = 4.27, p = 2.97×10⁻⁵), indicating that the predictive value of IMM-AGE is not reducible to pre-procedural cardiac status; and (iii) finally, this association survived a broader model incorporating pre-procedural laboratory, echocardiographic and clinical predictors, including biomarkers of myocardial stress, inflammation and fibrosis (**Supplementary Fig. 5H**; log-odds=8.06, p=2.8×10^⁻2^, **Supplementary Note 1**). Moreover, beyond recovery, pre-procedural IMM-AGE also stratified mortality risk: patients who died during follow-up had higher baseline IMM-AGE than survivors (**Fig. 4F**; Wilcoxon p = 3.5 × 10^−2^), further supporting the ability of IMM-AGE to resolve prognostic heterogeneity within this clinically vulnerable group.

### IMM-AGE distinguishes future cardiovascular events among clinically similar individuals

We next extended the evaluation of flow-based IMM-AGE to a second, complementary cardiovascular setting: prospective prediction of incident events. Even among individuals with similar conventional risk profiles, cardiovascular outcomes can diverge substantially, pointing to residual risk not captured by standard clinical assessment. Despite the known role of inflammation in cardiovascular disease, incorporation of immune-based information (e.g. CRP) into predictive models is rare, with no established way to capture a higher-level immune-state^7,29–31^. We therefore asked whether flow-based IMM-AGE could help distinguish future cardiovascular events among otherwise clinically similar individuals. Whereas the UKB provided population-scale context, the Project Baseline Health Study (PBHS) combined deep phenotyping and longitudinal follow-up with flow-cytometry data, enabling direct evaluation of flow-based IMM-AGE at the individual level. BHS comprised 2,234 adults aged 18– 92 years who underwent annual clinical assessment and were followed for a median of 1,463 days. During follow-up, 122 individuals experienced a first cardiovascular event, including coronary, cerebrovascular, thrombotic, arrhythmic, structural or fatal outcomes (**Fig. 5A; Supplementary Table 16**; see Methods). Of these, 105 cases and 949 controls had complete data across all relevant clinical, pre-existing risk factors and were retained for matched analysis.

**Figure 5.**
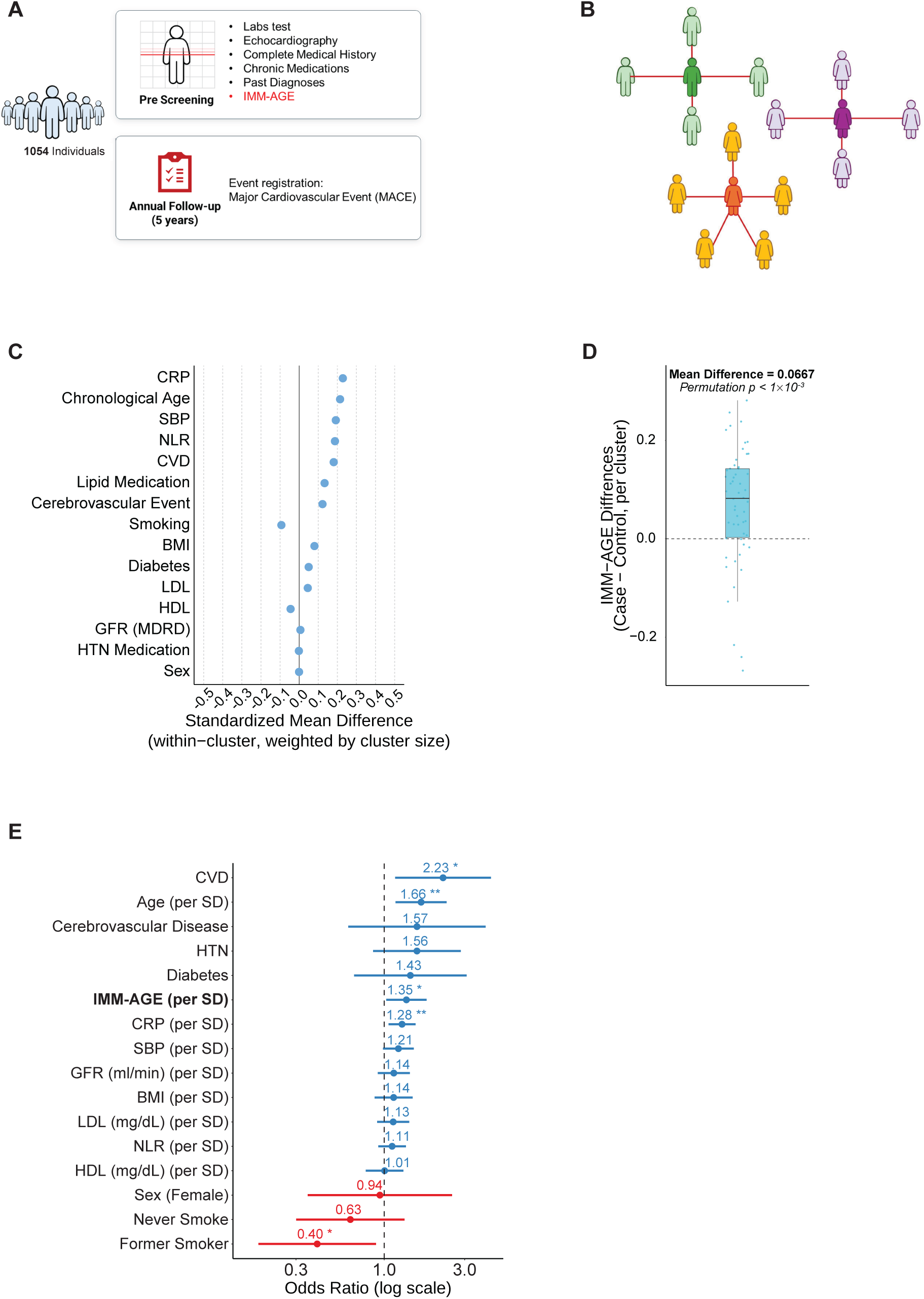
IMM-AGE independently predicts major adverse cardiovascular events among clinically matched individuals. **(A)** Retrospective study design and multidimensional phenotyping of the Project Baseline Health Study (BHS) cohort. Schematic illustrating the longitudinal study workflow for 2,234 participants, who were followed for a median of 1,463 days during which incident major adverse cardiovascular events were recorded (MACE; *n* = 105). **(B)** Network-based similarity matching of cardiovascular event cases and controls. Schematic illustrating the high-dimensional matching strategy used to identify clinically similar case-control groups based on 15 clinical and laboratory covariates. Each case was matched to its ten nearest controls, and overlapping matched sets were subsequently merged into shared network clusters. **(C)** Covariate balance across matched case-control clusters. Love plot showing cluster-aware standardized mean differences (SMDs) for 15 demographic, clinical, laboratory, and treatment variables across 54 merged case-control clusters. SMDs were calculated within each cluster and averaged across clusters, weighted by harmonic cluster size. Dashed vertical lines indicate the conventional threshold for covariate balance (SMD = ±0.1). **(D)** Higher baseline IMM-AGE precedes incident cardiovascular events in matched participants. Baseline IMM-AGE scores were compared between participants who subsequently experienced cardiovascular events and their clinically matched controls across matched clusters. The mean case-control difference was 0.0667, with significance assessed by permutation of case-control labels within clusters (1,000 permutations; *P* < 0.001). **(E)** Baseline IMM-AGE independently predicts major adverse cardiovascular events. Forest plot showing odds ratios from a mixed-effects logistic regression model for incident MACE, adjusted for 14 conventional cardiovascular risk factors with a random intercept for matched clusters. IMM-AGE remained independently associated with incident MACE (OR = 1.35; 95% CI, 1.02–1.78; *P* = 0.03). CVD, cardiovascular disease; HTN, hypertension defined by blood-pressure treatment; CRP, C-reactive protein; NLR, neutrophil-to-lymphocyte ratio; SBP, systolic blood pressure. For continuous variables, ORs are reported per 1-SD increase.

To generate highly matched case-control groups, we constructed a similarity network from demographic, clinical, laboratory and treatment variables, matched each case to one or more of its most similar controls, and merged overlapping clusters (**Fig. 5B**; **Supplementary Fig. 6A-D**; see Methods). Matching achieved adequate balance across 8 of the 15 covariates, and all covariates were therefore included as fixed effects in subsequent models to account for residual confounding (**Fig. 5C**, see Methods). Within this matched framework, individuals who subsequently experienced cardiovascular events had significantly higher IMM-AGE scores than their matched controls (**Fig. 5D**; mean difference = 0.0667 IMM-AGE units; permutation p < 1 × 10^−3^, 1,000 permutations). Importantly, this difference was specific to case-control contrasts, exceeding the IMM-AGE differences seen between matched controls within the same clusters (**Supplementary Fig. 6E**; permutation p < 1 × 10^−3^, 1,000 permutations).

We next asked whether IMM-AGE remained informative after accounting for conventional risk factors within the matched design. In a conditional logistic regression model stratified by merged clusters and adjusted for conventional risk factors, IMM-AGE remained significantly associated with cardiovascular event status (**Fig. 5F**; OR = 1.35, 95% CI 1.02-1.78; p=3×10^−2^; see Methods). However, a complementary Cox proportional hazard analysis did not reach statistical significance (**Extended Data Fig. 1**). Notably, a 0.06-unit difference in IMM-AGE between clinically similar individuals corresponded to an estimated ∼11% increase in the odds of a cardiovascular event. Chronological age, CRP, smoking status and history of CVD were also independently associated with event status (p=4.6×10^−3^, 9×10^−3^, p=2.5×10⁻², p=1.5×10⁻², respectively). Finally, inclusion of IMM-AGE significantly improved model fit relative to a reduced model containing conventional risk factors alone (likelihood ratio test, χ²(1) = 5.233, p=2.2×10^−2^), collectively indicating that IMM-AGE captures additional prospective cardiovascular risk among otherwise similar individuals.

## Discussion

Considering the central role immune processes play in cardiovascular injury, remodeling and disease progression, cardiovascular risk assessment does little to incorporate immune state, reflecting in part the lack of a quantitative and clinically tractable measure. Here, we establish that immune aging, as quantified by IMM-AGE, is a distinct and clinically relevant dimension of cardiovascular disease-risk. Across diverse cardiovascular settings, higher IMM-AGE tracked cardiovascular morbidity, prediction of future events, intervention outcomes, mortality, and provided prognostic information beyond conventional risk factors, CRP and current risk-assessment approaches. Importantly, when incorporated into PREVENT, the incremental value of IMM-AGE was not limited to modest gains in overall discrimination but translated into movement across clinically relevant risk categories. Moreover, we establish a reduced-marker flow-cytometry implementation of IMM-AGE compatible with standard clinical instruments, providing a practical path toward routine clinical use. Applied in prospective and interventional settings, flow-based IMM-AGE identified patients at risk of maladaptive recovery following TAVR and distinguished individuals who subsequently experienced cardiovascular events from clinically similar controls. Taken together, our findings both support and enable the incorporation of immune aging into precision cardiovascular risk assessment.

Recent work has established that complex immune states can be distilled into quantitative measures of aging and health. The inflammatory aging clock iAge links systemic inflammation to multimorbidity, frailty and vascular dysfunction^32^, while a unified immune-health metric distinguishes healthy from immune-mediated disease states across diverse conditions^33^. IMM-AGE extends this framework in a distinct direction. Rather than being trained against chronological age or a specific clinical endpoint, it was derived, in an unsupervised manner, from the natural, coordinated remodeling of immune-cell composition that occurs during immune-aging^19^. Its association with disease is therefore not a given, nor is its ability to generalize across cardiovascular settings. This distinction is particularly relevant given parallel advances in high-dimensional cardiovascular biomarker models, which can achieve stronger discrimination when trained for specific outcomes^17,18^. A supervised panel optimized for a single endpoint is expected to outperform a fixed, outcome-agnostic metric, such as IMM-AGE, on that same endpoint but does so at the expense of generalizability. By contrast, the outcome-agnostic nature of IMM-AGE supports broader applicability: the same underlying, fixed metric remained informative across diverse cardiovascular conditions, and across prospective and interventional settings, despite no cardiovascular outcome contributing to its original derivation. This allowed us to ask a different question: not whether a cardiovascular endpoint could be predicted *de novo*, but whether the process of immune aging is relevant to cardiovascular disease and whether it provides clinically meaningful information beyond established risk-assessment approaches.

For that breadth to translate into clinical utility, IMM-AGE must also be portable across platforms and, ultimately, measurable in routine clinical settings. We therefore developed reference-free, low-dimensional representations of IMM-AGE across molecular and cellular modalities, each serving a different translational purpose. The transcriptomic and proteomic representations extend IMM-AGE into existing research cohorts, allowing the biology and clinical correlates of immune aging to be interrogated at scale across diverse populations and datasets. Beyond cardiovascular disease, these representations may provide a means to identify additional clinical contexts in which immune aging may be relevant, thereby helping to prioritize settings for prospective evaluation. By contrast, the reduced-marker flow-cytometry implementation provides a more direct route toward clinical measurement. Whereas broad proteomic profiling requires specialized platforms and large analyte panels^17,18^, a defined immune signature measured by flow can be implemented using instrumentation already available in standard clinical laboratories. The potential value of such a readout was evident across prospective and interventional settings, particularly in TAVR, where existing clinical and machine-learning models predict hard outcomes poorly and standard surgical scores are inconsistent^26^, yet pre-existing inflammatory phenotypes have been linked to adverse remodelling and post-procedural mortality^27^. In this setting, pre-procedural IMM-AGE provided a quantitative, system-level readout of immune state that identified patients at risk of maladaptive early cardiac recovery. Notably, the optimized flow panels were composed predominantly of T-cell populations, yet retained the broader IMM-AGE signal originally defined across T-cell, B-cell and myeloid compartments. A handful of T-cell measurements thus provide a readable window into a system-level immune state, reflecting the coordinated correlation structure of immune-cell remodeling that is captured by IMM-AGE. Together, these complementary implementations establish a translational framework in which molecular representations enable broad discovery, while flow provides a clinically tractable route for prospective evaluation.

Moving IMM-AGE toward clinical use will require a shift from establishing prognostic relevance to clinical interpretation, standardization and prospective validation. Reference levels and age- and sex-specific distributions will be required to contextualize an individual’s IMM-AGE, alongside standardization of the reduced flow-cytometry panel across laboratories and prospective definition of thresholds that meaningfully inform clinical decisions. A central outstanding question is why immune aging is so closely linked to cardiovascular disease. Our findings are consistent with advanced immune aging preceding overt cardiovascular manifestations rather than simply reflecting established disease: in the UK Biobank, prior cardiovascular disease accounted for only a small fraction of IMM-AGE variance, while baseline IMM-AGE predicted subsequent cardiovascular events, and, independently, flow-based IMM-AGE distinguished individuals who subsequently experienced cardiovascular events from clinically similar controls in the Baseline Health Study. Determining which components of the immune-aging program contribute to this vulnerability, and whether that relationship is causal, will require dedicated mechanistic investigation. The prominence of T-cell populations in our reduced panel, together with growing evidence linking T-cell biology to cardiovascular disease^34,35^, provides a natural entry point for such mechanistic investigation, while the system-level nature of IMM-AGE argues against reducing the signal to any single lineage. Resolving these mechanisms will also begin to determine whether immune aging is itself modifiable and, ultimately, whether shifting IMM-AGE toward a younger immune state can reduce cardiovascular risk. Collectively, by converting a complex trajectory of immune aging into a portable and clinically measurable variable, this study provides a foundation not only for incorporating immune-system state into precision cardiovascular risk assessment, but also for testing whether immune aging can become a future target for cardiovascular prevention.

## Acknowledgements

This research was supported by NIH joint NIAID & NIA award P01AI153559, the ISRAEL SCIENCE FOUNDATION (grant No. 1626/20), within the Israel Precision Medicine Partnership program and the Impetus Longevity award.

## Author Contributions

Conceptualization: E.F., T.J.F.C., N.M. and S.S.S.O.

Methodology: E.F., T.J.F.C., N.M., S.S.S.O.

Software: E.F., N.G., E.J.S.

Validation: E.F., K.M., S.M., Y.K.

Formal analysis: E.F., E.J.S., N.G., M.L., S.Short, M.C.

Investigation: E.F.,, S.Short, M.C., S.S.S., B.C., S.B., M.L., O.C.

Data Generation: S.M., K.M., S.Short, M.C., S.S.S., X.J., Y.K.

Resources: K.W.M., W.F.F., B.K., F.H.

Data curation - E.F., E.J.S., B.C., S.B., K.M., S.M., S.Short, M.C., S.S.S.

Writing: E.F., T.J.F.C., N.M., F.H. and S.S.S.O.

Supervision: S.S.S.O., F.H., T.J.F.C., N.M., H.T.M., P.K.P.N., M.M.D.

Funding acquisition: S.S.S.O., M.M.D., S.Shah,, E.F.

All authors approved the final manuscript.

## Methods

### Human cohorts and sample sourcing

#### The Stanford-Ellison Longitudinal Aging (SELA) cohort

Briefly, SELA, is an ongoing longitudinal cohort in which peripheral blood is collected annually and deeply immunophenotyped. A previously published analysis of this cohort, encompassing the first nine-year period (2007-2015) identified an immune-aging trajectory and subsequently led to the development of IMM-AGE - the first high resolution metric of immune aging^19^. This cohort’s samples and measurements are referred to here, throughout, when describing SELA: 135 healthy adults spanning two age ranges at enrollment, including 63 young adults (20-31 years) and 72 older adults (60-96 years). Participants were assessed as healthy at enrollment based on medical history and vital signs, and were followed longitudinally with repeated sampling - with participants asked to return for regular, annual visits post enrollment. Peripheral blood (65 mL) was collected and processed into peripheral blood mononuclear cells (PBMCs) and serum using standardized procedures at the Stanford Human Immune Monitoring Center. Immune profiling included longitudinal cellular phenotyping (flow cytometry in initial years; mass cytometry (CyTOF) in proceding years), functional signaling responses to cytokine stimulation measured by phospho-flow assays, serum proteomics, and whole-blood gene expression profiling; complementary “snapshot” profiling was also performed on longitudinal samples from a subset of participants assayed together to minimize year-to-year technical variation. Samples were collected at the Stanford Clinical and Translational Research Unit in the context of an annual influenza-vaccination study (SLVP015; ClinicalTrials.gov NCT01827462)

#### Framingham Heart Study (FHS)

FHS is a community-based, multigenerational prospective cohort initiated in Framingham, Massachusetts, in 1948 to investigate determinants of cardiovascular disease. For the present analysis, we used participants from the Offspring cohort, which enrolled 5,124 adult children of the original FHS participants and their spouses beginning in 1971, with repeated standardized examinations performed approximately every 4-8 years and continuous surveillance for clinical outcomes. In line with the analyses we have performed by Alpert et al. to show IMM-AGE captures cardio and survival signal^19^, we perform an analysis based on Offspring participants with available whole-blood gene-expression profiling (n=2131), generated using the Affymetrix Human Exon 1.0 ST Array. Institutional Review Board (IRB) approval to analyze FHS data was obtained from the **Rambam ethics board** (IRB number RMB-D-0270-24).

#### UK Biobank (UKB) cohort

We identified participants with available plasma proteomic measurements in the UK Biobank Pharma Proteomics Project (n = 52995), and extracted their corresponding baseline clinical variables, self-reported diagnoses, medication data, and linked outcome records. This research was conducted using the UKB resource under an approved application (Application ID 1164153). We classified medications into blood-pressure-lowering and lipid-lowering therapies by matching all reported treatments to curated drug lists (**Supplementary Table 17**). Baseline clinical variables included age, sex, smoking status, body mass index, systolic blood pressure, lipids, glycated hemoglobin, and estimated glomerular filtration rate (eGFR), with eGFR calculated using the CKD-EPI equation. Prevalent conditions were defined from self-reported diagnoses. Type 2 diabetes and prevalent cardiovascular disease were assigned using curated diagnosis lists (**Supplementary Table 18**).In addition, comorbidities were summarized into clinically meaningful categories based on predefined groupings of diagnoses (**Supplementary Table 18**).

Incident cardiovascular outcomes were defined using curated UKB event fields corresponding to major atherosclerotic cardiovascular disease and heart failure (**Supplementary Table 5**). Events were required to occur after the baseline assessment-centre visit. A composite outcome indicator was defined as the occurrence of any qualifying event. Time-to-event was calculated from the assessment date to the first event. Participants without an event were censored at death or last recorded follow-up, whichever occurred first. Over a median follow-up of 13.5 years (IQR: 10.1– 14.7), n = 4978 participants experienced an incident cardiovascular event.

#### Transcatheter Aortic Valve Replacement (TAVR) cohort

Briefly, this cohort - the details of which have been previously published^28^ - is a prospective, single-center registry study of 112 adult patients with severe calcific aortic stenosis undergoing TAVR at Stanford University Medical Center. The mean age of the cohort was 84 years and 59% were male. At baseline, fasting blood samples were collected and stored at −80°C for centralized biomarker analysis, performed in a manner blind to clinical data. Plasma levels of B-type natriuretic peptide (BNP), high-sensitivity troponin I (hs-TnI), C-reactive protein (CRP), galectin-3 (GAL-3), growth differentiation factor-15 (GDF-15), and cystatin-C (Cys-C) were measured using standardized immunoassays. Comprehensive transthoracic echocardiography was performed at baseline and repeated at follow-up (1 month, and at 1 year in a subset of patients), including assessment of left ventricular ejection fraction (LVEF), global longitudinal strain (GLS), left ventricular mass index (LVMI), left atrial volume index (LAVI), and indices of aortic valve severity. Clinical risk was assessed using the Society of Thoracic Surgeons (STS) score and frailty measures, and participants were followed longitudinally for all-cause mortality and echocardiographic evidence of ventricular remodeling. The study was approved by the Stanford University Institutional Review Board (protocol 34394), and all participants provided written informed consent.

Echocardiography was acquired according to the ASE guideline recommendations using a standard echocardiographic machine (iE33, and EPIQ 7C; Philips Medical Imaging). Echocardiographic views were obtained in M-mode, two-dimensional (2D) and color tissue Doppler modes. The following measurements were performed: (i) left ventricular (LV) end-systolic and end-diastolic volumes; (ii) ejection fraction (LVEF) using Simpson’s biplane method; (iii) LV internal diameter; (iv) interventricular septal and posterior wall thicknesses at end-diastole from the 2D image; (v) diastolic parameters with transmitral pulse doppler velocities,tissue doppler velocities of the mitral annulus and E/e’ mean ratio; and, finally, (vi) stroke volume and LV strain using Lagrangian strain by manual tracing. The Lagrangian method consists of measuring the myocardial initial length in end-diastole (L0) and the final length in end-systole (L1) and calculating LV strain values as 100 × (L1 − L0)/L04. GLS represents the average values of longitudinal strain calculated from the apical 4-, 3-, and 2-chamber views.

To assess the severity of aortic valve stenosis, we measured aortic valve area (using the continuity equation), peak and mean systolic transaortic pressure gradients (using the simplified Bernoulli equation), the ratio of left ventricular outflow tract velocity of blood flow to the aortic valve velocity and the global LV afterload. Severe aortic stenosis was defined according to ASE guidelines as an aortic valve area (AVA) of ≤1.0 cm2 or indexed AVA (AVAI) of ≤0.6 cm2/m2 and/or mean systolic aortic gradient >40 mmHg or peak velocity across the aortic valve >4 m/sec. To identify severe aortic stenosis, for the patients with LV systolic dysfunction and low-flow, low-gradient AS, the severity of AS was diagnosed and confirmed by low-dose dobutamine stress echocardiography.

#### Project Baseline Health Study (PBHS)

The PBHS study is a multicentric prospective longitudinal study whose goal is, via longitudinal deep-phenotype profiling, to characterize diversity at the level of the population and from this provide biomarkers of disease-related transitions. This study included participants from across various health spectrums from healthy participants, patients at risk of disease to participants with active disease. Participants were recruited across 4 centers in the United States, at Stanford University (Stanford, California), Duke University (Durham, North Carolina), and the California Health and Longevity Institute (Westlake Village, California) through a variety of methods such as IRB-approved advertisements, registries, care provider recommendation, and community events. Eligibility criteria required individuals to be older than 18 years old, a US resident, able to speak and read English and willing and able to comply with all aspects of the protocol. Exclusion criteria included sponsor employees and individuals working on Project Baseline and those with known, severe allergies to nickel or metal jewelry. Institutional Review Boards at Duke University and Stanford University approved the study protocol. A written informed consent was also signed by all participants. ClinicalTrials.gov Identifier: NCT03154346.

PBHS participants have been evaluated serially using clinical, molecular, imaging, cardiovascular imaging, sensor, self-reported, behavioral, psychological, environmental, and other health-related measurements. The measurements were performed at the start visit and at follow-up, which was performed annually for a total duration of 5 years. The PBHS study design has been previously published in detail^36^. In total, 2502 participants were enrolled.

### Experimental procedures

#### SELA Gene expression data generation for learning a cross-platform IMM-AGE signature

##### RNA extraction

RNA collection and extraction: Whole blood (WB) (PAXgene) was collected directly into PAXgene Blood RNA Tubes (Cat. # 762165, Waters Biosciences). Samples were incubated in collection tubes at room temperature then stored at −80C as per the manufacturer’s instructions within 4h. Total RNA was isolated according to the manufacturer’s protocol by using a PAXgene RNA blood kit (Cat, #762164, Qiagen), carried out at room temperature by a QIAcube automated robot (QIAcube). PBMCs were processed from peripheral blood by standard procedures and stored in liquid nitrogen until use. Cell pellets were resuspended in the RLT buffer (QIAGEN) with 1% 2-mercaptoethanol (BME) then RNeasy mini kit (Cat. #74104, QIAGEN) was used for total RNA extraction from PBMC on QIAcube automation. Total RNA yield was assessed using the Thermo Scientific NanoDrop 1000 micro-volume spectrophotometer (using absorbance at 260 nm, and the ratio of 260/280 and 260/230). RNA integrity was assessed using Agilent’s Bioanalyzer NANO Lab-on-Chip instrument (Agilent).

##### Microarray processing and analysis

Affymetrix GeneChip® PrimeView™ Human Gene Expression microarrays were used according to the manufacturer’s protocol. Biotin-labeled antisense complementary RNA (cRNA) was synthesized and hybridized using the GeneChip® 3′ IVT Express Kit, starting from 50 ng of total RNA. Briefly, total RNA underwent reverse transcription to generate first-strand complementary DNA (cDNA), followed by second-strand synthesis to produce double-stranded cDNA templates. *in-vitro* transcription was then performed to generate biotin-labeled amplified RNA (aRNA), which was subsequently purified and fragmented to prepare samples for hybridization onto GeneChip® 3′ expression arrays. Arrays were scanned using the Affymetrix GeneChip® Scanner, and data were extracted and subjected to preliminary analysis using the GeneChip® PrimeView™ Human Gene Expression Assay Software Module v1.0. Assay data were generated and reported in multiple formats, including TIFF image files, .DAT, .CEL, .CHP, and associated metadata files (e.g., .ARR, MD5), for downstream analysis.WB and PBMCs gene expression profiles were generated using Affymetrix PrimeView microarrays. Raw .CEL files were imported using the *ReadAffy* function from the *affy* R package (v1.78.0, Gautier et al. 2004^37^). Probe-level preprocessing was performed using the *expresso* function from the *affy* package, with the following parameters: background correction using Robust Multi-array Average (RMA; bgcorrect.method = “rma”), quantile normalization (normalize.method = “quantiles”), perfect match probe correction (pmcorrect.method = “pmonly”), and summarization via median polish (summary.method = “medianpolish”).

Probe-to-gene annotation was carried out using the *select* function from the *AnnotationDbi* package (v1.64.1, Pagès et al. 2023), with annotation data provided via a custom SQLite database created using AnnotationForge (v1.40.0, Carlson et al. 2026) and the Affymetrix PrimeView.na36 annotation file. The resulting gene-level expression matrix was generated by mapping probes to gene symbols. For genes represented by a single probe, the probe intensity was used directly. For genes with multiple probes, expression values were aggregated by computing the median across probes (*aggregate* function from *stats* package, base R). Probes without valid gene symbols or with missing values were excluded from all downstream analyses.

#### RNA sequencing

RNA sequencing libraries were prepared using the KAPA mRNA HyperPrep Kit (Roche; KK8580) in combination with the IDT for Illumina® Dual Index Adapter Kit (Illumina; Cat. #20021454), according to the manufacturer’s protocols. Briefly, polyadenylated mRNA was enriched from total RNA using magnetic oligo-dT beads. Captured mRNA was fragmented using heat and magnesium, followed by first-strand cDNA synthesis using random priming. Second-strand cDNA synthesis incorporating dUTP, A-tailing, adapter ligation, library amplification, and KAPA Pure Beads clean-up steps were performed to generate sequencing libraries. Incorporation of dUTP during second-strand synthesis enabled strand-specific sequencing, as the dUTP-marked strand was not amplified during library amplification.

Final libraries were assessed for quality and fragment size distribution using the Agilent 2100 Bioanalyzer with the High Sensitivity DNA Kit. Equimolar amounts of each library were pooled and sequenced on an Illumina NovaSeq 6000 platform using an S4 flow cell. FASTQ files were generated using bcl2fastq2 Conversion Software v2.19 (Illumina). Sequencing reads were aligned to the human reference genome (GRCh38) using STAR (v2.7.10a; Dobin et al., 2013), with gene-level read counts quantified in GeneCounts mode. The genome index was generated using the GENCODE v43 primary assembly annotation. For samples sequenced across multiple lanes, FASTQ files from the same sample were concatenated prior to alignment. STAR output files (ReadsPerGene.out.tab) were collected for downstream analysis; strand-specific counts were used.

#### RNAseq data pre-processing

Gene-level raw counts were imported into R using the data.table package (v1.14.10, Barrett et al. 2024). Sample metadata were used to retain only PBMC-derived samples. Low-expression genes were filtered using *filterByExpr* function from the *edgeR* package (v3.42.4; Robinson et al., 2010) (minimum count threshold of 10), and library sizes were recalculated. Trimmed mean of M-values (TMM) normalization was applied using *calcNormFactors*, and normalized expression values were computed as counts per million (CPM) using the *edgeR* package (v3.42.4; Robinson et al., 2010) and *limma* (v3.56.2; Ritchie et al., 2015).

Gene annotation was performed using the *biomaRt* package (v2.58.0; Durinck et al., 2009) by querying the hsapiens_gene_ensembl dataset from Ensembl. Ensembl gene IDs were mapped to HGNC gene symbols using *getBM*. Genes mapping to a unique HGNC symbol were retained directly. For genes with multiple Ensembl IDs mapping to the same HGNC symbol, CPM values were aggregated by taking the median across transcripts. Genes lacking an HGNC symbol were retained under their Ensembl gene ID.

#### Flow cytometry immunophenotyping

PBMC from the SELA cohort (**Supplementary Table 11**) and TAVR cohort (**Supplementary Table 12**) were thawed in warm culture medium, washed twice, and resuspended at a concentration of 1 × 10⁶ viable cells/mL. Cells were plated in 96-well deep-well plates at 0.5 × 10⁶ cells per well in 100 µL. For viability assessment, cells were stained with a live/dead dye for 15 min at room temperature, followed by washing with FACS buffer (PBS supplemented with 2% fetal bovine serum and 0.1% sodium azide). Cells were then incubated with an antibody cocktail (**Supplementary Table 19**) for 30 min at room temperature. After staining, cells were washed three times with FACS buffer and resuspended in 200 µL FACS buffer for acquisition.

Data were acquired on an LSRII flow cytometer (BD Biosciences) using DIVA 6.0 software, with 200,000 lymphocytes collected per sample. Flow cytometry data were analyzed using FlowJo v9.3. Gating was performed by first selecting live cells based on forward and side scatter properties, followed by exclusion of doublets using forward scatter area versus height, and subsequent identification of cell subsets according to established gating strategies (**Supplementary Fig. 3D**).

#### Computational analysis

All statistical analyses were conducted in R (version 4.3.3)

### Methods for IMM-AGE estimation

#### Gene-ratio based IMM-AGE assessment

##### Universal gene-expression-based signature for IMM-AGE estimation

To identify gene expression ratios required for IMM-AGE estimation, we integrated cytometry and gene expression data from whole blood profiled in the SELA cohort. Gene expression data (Affymetrix) and cytometry features were preprocessed using R and Bioconductor packages: *GEOquery* (v2.66.0, Davis et al. 2007^38^) and *GSVA* (v1.50.0, Hänzelmann et al. 2013^39^).

Expression data were filtered by removing genes with low median intensity across individuals. The cutoff (median intensity = 4.2) was chosen empirically from the bimodal distribution of probe intensities, corresponding to the transition from background-level to expressed signal. Genes were retained only if expression exceeded this threshold in at least as many samples as the smaller age group, ensuring robust detection across both young and old individuals. To determine each gene’s individual association with IMM-AGE, we, for each year (2012-2015), computed Spearman correlations between filtered gene expression values and cytometry-defined IMM-AGE features (*cor.test*, method = “spearman”). Gene-cell pairs with consistent correlation direction across all years and an average correlation strength of |r|≥ 0.4 were retained (150 out of 14,956 genes). Pairwise gene expression ratios were subsequently calculated between all retained genes. Of 11,175 total gene ratios, 8,117 exhibited consistent correlation with IMM-AGE scores across years, and of those, 1,088 displayed an adjusted combined p-value < 0.01 and a mean Spearman correlation larger than 0.6. P-values were combined across years using Fisher’s method from the *metap* package (v1.9, Dewey et al. 2025^40^). Ratios were further prioritized and selected using stability selected LASSO regression (ssLASSO) (*stabsel* from the *stabs* package, v0.6-4, Hofner et al. 2015^41^), retaining only ratios appearing in at least two years with a selection probability ≥ 0.05. Of 1,088 gene ratios, 163 were selected by the regression model to build the final signature and comprised information from 105 unique genes (**Supplementary Table 20**).

Parallel processing (*foreach* v1.5.2, Microsoft, Weston et al. 2022; and *doParallel* v1.0.17 Microsoft, Weston et al. 2022) was employed throughout for computational efficiency using up to 35 cores. Visualization of correlation results was performed using *ggplot2* (v4.0.1, Wickham et al. 2016^42^, and *ggpubr* (v0.6.0, Kassambara et al. 2026).

##### Validation of the ratio-based IMM-AGE signature in SELA

To validate our novel gene ratio-based transcriptomic IMM-AGE signature, we applied it to SELA samples - with previously computed IMM-AGE scores derived from CyTOF data-using single-sample gene set enrichment analysis (ssGSEA) via the *GSVA* (v1.50.0, Hänzelmann et al. 2013) (**Supplementary Table 1**). Briefly, for each sample, the IMM-AGE score was defined as the difference between the enrichment scores of the up- and down-regulated ratio sets. Resulting transcriptomic IMM-AGE scores were compared to CyTOF-based IMM-AGE estimates using Spearman correlation.

#### Proteomic based IMM-AGE assessment

##### Feature selection and signature training

To enable application of IMM-AGE across proteomic platforms, we first restricted the candidate feature space to proteins with evidence of cross-platform stability across 9 studies (**Supplementary table 21**). Specifically, proteins were retained if they showed high inter-platform agreement between SomaLogic and Olink measurements, defined as Spearman’s rank correlation > 0.7 observed in at least 2 independent studies, or Spearman’s rank correlation > 0.6 with at least one cis pQTL evident. This filtering step was used to prioritize proteins whose measured abundance patterns were concordant across assay technologies, thereby improving transferability from SomaScan-based discovery data to Olink-based external cohorts.

To derive the proteomic IMM-AGE signature, we trained the model on a discovery cohort from the SELA study, comprising 70 samples collected from 40 individuals. High-dimensional proteomic profiling was performed using the SomaScan® 11K Assay, for which ground-truth IMM-AGE scores had been previously established. SomaScan RFU values were log-transformed to better align their scale with Olink NPX values, and each protein was standardized within cohort prior to feature selection. Additional feature selection was performed on this standardized protein matrix to identify proteins associated with IMM-AGE. The final signature comprised 145 proteins, partitioned into modules positively associated with IMM-AGE (UP, n = 102) and negatively associated with IMM-AGE (DOWN, n = 43). This UP–DOWN structure was then used to estimate IMM-AGE as a relative proteomic score in external cohorts (**Supplementary Table 3**).

##### Data Harmonization and Preprocessing

To address platform-specific technical variance, dynamic range differences, and chemical amplification properties across diverse proteomic technologies, we implemented an internal standardization protocol for all datasets. Prior to analysis, raw SomaScan relative fluorescent units (RFU) underwent a log⁡(x+1) transformation to harmonize the data structure with Olink’s inherently log2 scaled Normalized Protein eXpression (NPX) data. After log transformation, protein levels were standardized within each cohort by converting them to Z-scores (mean = 0, standard deviation = 1).

##### IMM-AGE scoring formulation

To apply the signature to external datasets, including the UKB Olink proteomics cohort, protein values were first standardized within each cohort. IMM-AGE was then calculated by aggregating the cohort-specific Z-scores of the signature proteins. The final score was defined as the difference between the summed signal of proteins positively associated with IMM-AGE and the summed signal of proteins negatively associated with IMM-AGE, according to the following equation:

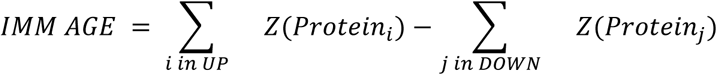

#### Design and Evaluation of Reduced-Marker IMM-AGE Panels for Clinical Flow Cytometry

##### Sample collection

Mass spectrometry (CyTOF) immune profiling data was previously published in Alpert et al. Data was obtained from participants in the SELA cohort. For the present analysis, we used CyTOF data from a subset of 262 individuals profiled between 2012 and 2015. Immune cell frequencies derived from these data were used to compute IMM-AGE scores and assess the performance of reduced-marker cytometry panels.

##### Panel construction

The original IMM-AGE design involved simultaneous measurement of 18 immune cell types, often requiring evaluation of approximately 25 cell surface markers via CyTOF. To enable clinical application of IMM-AGE, we sought to design flow cytometry panels compatible with standard cytometers, which typically support no more than 8 markers. We systematically enumerated all possible, viable 8-marker combinations from the full IMM-AGE panel. Panels were retained if the selected markers enabled complete phenotyping of at least three predefined immune cell subsets, based on established gating definitions. This filtering process yielded 255 valid marker combinations.

##### Projection on simulated panel

To enable single-sample estimation of IMM-AGE from reduced-marker flow cytometry data, we developed a projection pipeline based on diffusion map embedding. The original IMM-AGE trajectory was derived from CyTOF-based immune profiling of the SELA cohort and computed using the *DiffusionMap* function from the destiny package (v 3.18, Angerer et al; 2016^43^). This reference trajectory served as the embedding space for downstream sample projection.

New samples acquired using reduced 8-marker flow panels were first scaled to match the training data distribution. Projection of these samples into the diffusion map space was performed using the *dm_predict* function from *destiny*, with Minkowski distance (p = 1.5) as the similarity metric.

The resulting DC1 and DC2 coordinates were combined with the original CyTOF-based coordinates to generate a comparative trajectory.

To assign IMM-AGE scores to the projected samples, we used a nearest-neighbor averaging approach. Briefly, for each new sample, the three nearest original SELA samples were identified in DC1-DC2 space using the *nn2* function from the *RANN* package (v2.6.1, Jefferis et al. 2024). The resulting IMM-AGE scores (pred_pt) provided a quantitative placement of each flow cytometry sample along the reference immune aging trajectory.

##### Trajectory concordance analysis

Using CyTOF data from the SELA cohort, we simulated cell subset frequencies for each valid 8-marker panel and used them to derive IMM-AGE scores per individual by assembling a trajectory which is based on the longitudinal change in cell frequencies. These scores were then ordered to form sample-level immune aging trajectories. To evaluate the extent to which reduced-marker panels preserved the biological signal of the full IMM-AGE panel, we calculated Kendall’s tau correlation between the sample rankings from each reduced panel and those obtained using the full panel.

##### Prognostic performance assessment

The clinical relevance of each reduced-marker panel was evaluated by testing whether simplified panels preserved the prognostic signal previously reported for the full IMM-AGE framework in predicting all-cause mortality. Immune cell subsets from each panel were mapped to corresponding gene expression signatures by identifying genes whose expression levels correlate with the abundance of the respective immune populations. These signatures were applied to whole-blood gene expression profiles from participants in the Framingham Heart Study (n = 2,131), with follow-up for all-cause mortality (297 deaths over a maximum of 2,500 days). Cox proportional hazards regression was performed to assess associations with mortality, adjusting IMM-AGE for age, sex, smoking status, total cholesterol, CMV serostatus, blood-pressure category, diabetes status, and prior cardiovascular disease. Predictive performance was quantified using the concordance index (C-index).

##### Selection of candidate panels

Panels ranking in the top 10% for both trajectory concordance (Kendall’s tau) and prognostic performance (C-index) were selected as high-performing candidates. From this subset, three representative 8-marker panels were selected and finalized for testing.

##### Selection and preprocessing of immune features for IMM-AGE flow estimation for panel validation

To facilitate IMM-AGE assessment on a clinically compatible flow-cytometer we characterized immune aging across samples by, we selected a panel of seven cytometry-derived gated immune cell populations previously identified as informative and sufficient for IMM-AGE dynamical characterization: total T cells, effector memory CD4⁺ T cells, naïve CD4⁺ T cells, effector CD8⁺ T cells, naïve CD8⁺ T cells, effector memory CD8⁺ T cells, and CD28⁻ CD8⁺ T cells.

All cytometry data were obtained from SELA cohort samples and read using readRDS() from an internal repository. After removing incomplete cases (complete.cases()), the data were scaled per feature using a robust z-score normalization within the 10th to 90th percentile range to suppress outlier influence.

##### Construction of the reference immune aging trajectory (Diffusion Map)

To learn the intrinsic structure of immune variation, we used the DiffusionMap() function from the destiny package (v 3.18, Angerer et al; 2016). A preprocessed and scaled SELA data matrix (features 46 samples) was embedded into a low-dimensional space using diffusion map and diffusion pseudotime (DPT) values were computed using DPT() from the same package. These DPT scores served as the IMM-AGE scores for the reference cohort.

### Computational analyses of cohorts

#### FHS cohort

##### Correlation of gene-ratio–based and original transcriptomic IMM-AGE scores

Gene-ratio-based IMM-AGE scores were generated to assess concordance with the original transcriptomic IMM-AGE scores. Predefined gene-expression ratios were computed from the filtered expression matrix, and single-sample enrichment scores were calculated using the *ssGSEA* function (GSVA R package, v1.50.5). The final ratio-based IMM-AGE score was defined as the difference between enrichment of ratios positively and negatively associated with IMM-AGE, then direction-aligned and standardized relative to age. Agreement with the original transcriptomic IMM-AGE score was assessed by Spearman correlation analysis.

##### Survival analysis

Survival analyses were performed largely as described previously in Alpert et al., with the key modification that gene-ratio–based IMM-AGE scores were used in place of the original transcriptomic IMM-AGE metric. In the Framingham offspring cohort, all-cause mortality was analyzed using multivariable Cox proportional hazards models implemented in the survival R package (v3.5-8, Therneau et al; 2001^44^), adjusting for age, sex, smoking, total cholesterol, CMV serostatus, blood-pressure class, diabetes status and prior cardiovascular disease. For Kaplan-Meier analysis, an adjusted IMM-AGE score was first obtained by regressing the ratio-based IMM-AGE score on the same risk factors using lasso regression with glmnet, and individuals were then stratified into high- and low-IMM-AGE groups based on the 60th percentile of the adjusted score. Follow-up time was truncated at 2,500 days, and survival curves were estimated with survival and visualized using ggplot2 (v4.0.1).

### IMM-AGE assessment across publicly available cardiovascular gene expression datasets

#### Data Acquisition and Preprocessing

We analyzed ten independent gene expression datasets obtained from the NCBI Gene Expression Omnibus (GEO), spanning a variety of transcriptomic platforms (Agilent, Affymetrix, and Illumina microarrays, and RNA sequencing). The cohorts included were: GSE20680, GSE90074, GSE180081, GSE218474, GSE59867, GSE58294, GSE209567, GSE38267 and GSE33463, along with the paired dataset, GSE20681.

Data retrieval was performed using the GEOquery package (v2.70.0) in R (v4.3.3). Unless stated otherwise below, each dataset was taken from its GEO series matrix in the form the submitters deposited it: a normalised, log2-scale expression matrix, used as supplied without further transformation. Where required, probe-level expression values were mapped to gene symbols. Probes lacking valid annotations were excluded. To address multiplicity (multiple probes mapping to the same gene), expression was summarized using the collapseRows function from the WGCNA package (v1.74) with the ‘MaxMean’ method, which retains the probe with the highest mean expression across samples while maintaining connectivity. As every feature in the signature is a ratio of two genes measured in the same sample, any scaling applied uniformly to a whole array (e.g. library size, scanner gain, per-array scaling) appears in both numerator and denominator and is there internally-normalized and insensitive to such transformations. Specifically, this covers the per-array scaling that several datasets applied (GSE20680 and GSE20681 scaled to a trimmed mean of 100, GSE33463 to a median of 256, and GSE58294 normalised to internal reference genes), none of which required adjustment. Four datasets are not distributed in this default form and were handled individually. GSE38267 is distributed as linear feature intensities rather than log2 values, and was log2-transformed before scoring. GSE180081 and GSE218474 carry no expression values in their series matrices, and relevant values were instead taken from supplementary files and handled according to the quantity supplied: raw counts (GSE218474) were converted to counts per million and log2-transformed as log2(CPM + 1), while RPKM values (GSE180081) were log2-transformed as log2(RPKM + 1) without a floor, since values below unity represent genuinely low expression rather than background. GSE90074, a two-colour common-reference design, was rebuilt from its sample channel with between-array quantile normalisation, as the per-gene reference term does not cancel within a gene-pair ratio. Spot filtering, missing value handling and probe-to-symbol mapping are given in full in the deposited script (GSE90074_Rebuild_From_Raw.R).

One residual per-gene term is disclosed rather than corrected. Data integrity was verified in every cohort by the within-sample rank percentile of the housekeeping genes ACTB, GAPDH and B2M, which ranged from 98.1 to 99.8 across the ten datasets (compared with 67.1 for GSE90074 before rebuilding).

#### IMM-AGE score computation

IMM-AGE scores were calculated using a ratio-based single-sample gene set enrichment analysis (ssGSEA, implemented via GSVA R package v1.50.5) using the aforementioned gene-ratio based IMM-AGE signature.

#### Case and control definitions

Cases and controls were defined from each series’ own phenotypic annotation. In GSE20681, the 99 matched pairs were taken from the “disease state” field (Case vs Control). In GSE20680, cases were samples annotated “Case”, denoting significant coronary stenosis (n = 87), and controls annotated “Control” (n = 52); the 56 samples annotated Intermediate were excluded, as this series does not resolve them to either arm. In GSE90074, cases were samples with obstructive coronary artery disease on angiography (n = 93) and controls those without (n = 50). In GSE180081, all participants presented for elective coronary catheterization; cases were those with coronary artery disease on invasive angiography, defined by the series as stenosis greater than 20% (recorded as “MID+”, angiographic grades D–F; n = 48), and controls as those below that threshold (“LOW”, grades A–B; n = 48). In GSE58294, cases were patients with cardioembolic stroke sampled at the earliest available timepoint, within three hours of the event and before thrombolytic treatment (n = 23), and controls were the series’ own non-stroke controls (n = 23); the 5-hour and 24-hour draws from the same patients were excluded so that each individual contributes one sample. In GSE59867, analysis was restricted to blood drawn on the first day of myocardial infarction at admission; cases were patients who subsequently progressed to heart failure (n = 9) and controls those who did not (n = 8), with the later timepoints and the stable coronary artery disease arm excluded. In GSE218474, cases were patients with aortic valve sclerosis (n = 44) and controls those without (n = 66), all presenting with acute coronary syndrome. In GSE209567, cases were Behçet patients of the vascular subtype (n = 9) and controls were healthy participants (n = 15); the mucocutaneous, ocular and complex subtypes were analysed separately. In GSE38267, cases were patients with pulmonary arterial hypertension (n = 13) and controls were healthy participants (n = 28), with the cystic fibrosis arm excluded. In GSE33463, cases were patients with idiopathic pulmonary arterial hypertension (n = 30) and controls were healthy participants (n = 41); the scleroderma and interstitial lung disease arms were not used in this contrast.

#### Effect-size estimation and significance testing

To enable comparable effect-size estimation across datasets and platforms, immune-aging differences were quantified as Hedges’ *g* - the difference in mean IMM-AGE between cases and controls divided by the pooled standard deviation, with a small-sample correction applied. Standardizing by the pooled standard deviation places cohorts of differing dynamic range on a common axis without rescaling the score itself. 95% confidence intervals were estimated using a t-based interval on the effect size. Uncorrected Cohen’s d and Cliff’s delta are reported alongside as sensitivity measures (**Supplementary Table 2**). Statistical significance was evaluated by label permutation testing on the difference in group means: case and control labels were reassigned at random with the group sizes held fixed, 1,000 times, and the two-sided p-values taken as the proportion of permutations whose absolute difference in means equalled or exceeded the observed one. The identical procedure was applied to every cohort, including those small enough for the permutation space to be enumerated exhaustively. All permutations used a fixed random seed (set.seed(100)) under R’s default Mersenne-Twister generator with rejection sampling; at 1,000 permutations the Monte Carlo standard error on a p-value near 0.05 is approximately 0.007. One cohorts (GSE218474) produced no permutations as extreme as the observed difference; for these the p-value is reported as below the resolution of the test, p < 1×10⁻³, rather than as zero. P-values were adjusted across the nine unpaired datasets using the Benjamini-Hochberg false-discovery-rate procedure. For the matched case-control cohort GSE20681, in which samples were paired by age and sex, a paired analysis was performed on the IMM-AGE scores of the 99 complete pairs. Within-pair differences were computed, and the effect was summarized as the mean within-pair difference and the paired Cohen’s d (d_z, the mean within-pair difference divided by the standard deviation of those differences), each with a 95% confidence interval. Significance was assessed by a two-sided paired sign-flip permutation test (1,000 permutations at the same fixed seed, randomly flipping the sign of each within-pair difference under the null of symmetry about zero), with the p-value computed as (1 + r)/(B + 1), where r is the number of permutations at least as extreme as the observed mean. Analyses were performed in R v4.3.3 with GEOquery v2.70.0, WGCNA v1.74, GSVA v1.50.5, readxl v1.4.3, effsize v0.8.1 and ggplot2 v4.0.2; the analysis scripts report these versions at run time.

### UKB cohort

#### IMM-AGE comparison between event and non-event groups

Participants were restricted to those aged 30–79 years, within predefined physiological ranges for key clinical variables, free of prevalent cardiovascular disease at baseline, with a classifiable current-smoking status, and with available IMM-AGE measurements, yielding n = 30,700 individuals. Analyses in this section were further restricted to individuals with complete data for all adjusted covariates (age, sex, glycated hemoglobin, body mass index, current smoking, systolic blood pressure, hypertension, dyslipidemia, type 2 diabetes, estimated glomerular filtration rate, HDL cholesterol, and total cholesterol), yielding a final analytical cohort of n = 22,024 individuals (Cases n = 2,760). Unadjusted differences in IMM-AGE between event and non-event groups were assessed using two-sided t-tests, and effect sizes were summarized using Cohen’s d.

To evaluate whether this difference persisted beyond conventional cardiovascular risk factors, IMM-AGE was regressed on baseline covariates including sex, age, glycated hemoglobin, body mass index, current smoking, systolic blood pressure, hypertension as defined by blood-pressure treatment, dyslipidemia as defined by lipid-lowering treatment, type 2 diabetes, estimated glomerular filtration rate, HDL cholesterol, and total cholesterol, using ridge regression (glmnet) with the regularization penalty selected by cross-validation. Covariate-adjusted IMM-AGE values were defined as residuals from this model, and residual distributions were then compared between event and non-event groups using two-sided t-tests.

#### Association of IMM-AGE with incident cardiovascular events

The association between IMM-AGE and incident major ASCVD/heart-failure events was evaluated using Cox proportional hazards regression using the *survival* R package (v3.5-8, Therneau et al; 2001^44^), with follow-up time defined from the baseline assessment-centre visit to the first qualifying event or censoring. The primary multivariable model included IMM-AGE together with the same covariate set used in the adjusted IMM-AGE comparison above (sex, age, glycated hemoglobin, body mass index, current smoking, systolic blood pressure, hypertension, dyslipidemia, type 2 diabetes, estimated glomerular filtration rate, HDL cholesterol, and total cholesterol). Continuous predictors were standardized before model fitting so that effect estimates could be interpreted per standard deviation increase. Hazard ratios and 95% confidence intervals were extracted from the fitted model. The proportional hazards assumption was assessed using Schoenfeld residuals. Schoenfeld residuals indicated no violation of the proportional hazards assumption for IMM-AGE (P = 0.4). Chronological age, by contrast, showed the expected time-dependent effect in cardiovascular risk26 (p=3.2×10-7). Modeling age as a time-varying covariate did not materially change the IMM-AGE estimate.

#### Variance decomposition of IMM-AGE

To quantify the contribution of baseline clinical and biological factors to variability in IMM-AGE, we performed variance decomposition using multivariable linear regression with IMM-AGE as the dependent variable. Candidate predictors included demographic variables, inflammatory markers, metabolic and lifestyle-related factors, treatment variables, and baseline disease burden indicators. Neutrophil-to-lymphocyte ratio was calculated as the neutrophil count divided by the lymphocyte count. Cardiovascular and non-cardiovascular morbidity were summarized as composite binary variables indicating the presence of at least one condition within each category.

Missing covariate data were handled using multiple imputation by chained equations. Imputation was performed using the mice R package (V3.19.0, Buuren & Groothuis-Oudshoorn; 2011^45^) with one imputed dataset (m = 1) and 10 iterations. Variable types were handled according to the imputation method assigned by mice, with predictive mean matching for continuous variables and logistic or polytomous regression for categorical variables as appropriate. The completed dataset was used for downstream analyses.

The primary model included age, sex, neutrophil-to-lymphocyte ratio, total cholesterol, estimated glomerular filtration rate, hypertension as defined by blood-pressure treatment, dyslipidemia as defined by lipid-lowering treatment, C-reactive protein, body mass index, current smoking, type 2 diabetes, cytomegalovirus seropositivity, cardiovascular disease burden, and non-cardiovascular disease burden. The proportion of variance explained was quantified using the coefficient of determination (R²). Relative contributions of individual predictors were estimated using the Lindeman-Merenda-Gold^46^ (LMG) decomposition using relaimpo R package (V2.2-7, Groemping; 2007^47^), and were further aggregated into broader domains.

#### Model fit comparison across nested COX models

Four nested Cox proportional-hazards models were fitted to evaluate the incremental predictive value of IMM-AGE and CRP beyond the AHA PREVENT score. All models used the PREVENT-predicted logit as a fixed offset (coefficient constrained to 1.0), such that added covariates test residual risk not captured by PREVENT. Models were: M0 (PREVENT only), M1 (PREVENT + CRP), M2 (PREVENT + IMM-AGE), and M3 (PREVENT + CRP + IMM-AGE). Akaike Information Criterion (AIC) was computed for each model overall and stratified by sex, with lower AIC indicating better model fit penalised for complexity.

#### Incremental discrimination by IMM-AGE and CRP

Harrell’s C-index was estimated for each nested model using Cox partial likelihood. To quantify the gain in discrimination attributable to each biomarker, the delta C-index (improvement over M0) was computed for each model. Because the C-index difference has no closed-form sampling distribution, we derived 95% confidence intervals from 1,000 bootstrap iterations, refitting both models on each resample and taking the 2.5th and 97.5th percentiles of the delta C-index distribution. We present both discrimination metrics overall and stratified by sex.

#### Time-dependent discrimination

To assess discrimination at the clinically relevant 10-year horizon, we estimated cumulative-sensitivity/dynamic-specificity time-dependent ROC curves for each nested model using the timeROC R package (v0.4; Blanche et al.^48^), with inverse-probability-of-censoring weighting to accommodate right-censored follow-up. For each model we used the predicted 10-year risk as the marker and evaluated the area under the time-dependent ROC curve (AUC) at 10 years. We obtained 95% confidence intervals from the influence-function–based variance estimator implemented in timeROC, and we compared time-dependent AUC between nested models using the package’s asymptotic test for paired AUC differences (the inverse-probability-of-censoring-weighted analogue of DeLong’s test). We report time-dependent AUC at 10 years and Harrell’s C-index separately, as they summarize discrimination at a single horizon versus global concordance across all event times, respectively, and are therefore not expected to coincide numerically.

#### Directional reclassification breakdown by event status

To clarify the interpretation of the continuous NRI, the underlying directional probabilities were visualised separately for events and non-events. For events, the proportion reclassified upward (correct) and downward (incorrect) by adding IMM-AGE to PREVENT are shown; NRI+ equals the difference between these two proportions. For non-events, downward reclassification is correct and upward is incorrect; NRI− equals their difference. The net NRI label in each panel makes explicit that NRI does not represent the proportion incorrectly reclassified, but rather the net directional balance.

#### PREVENT risk estimation and net reclassification improvement

For the PREVENT-based reclassification analyses, follow-up was administratively censored at 10 years: participants with an event after 3,650 days were reclassified as event-free for this horizon-specific analysis. After merging IMM-AGE with the PREVENT-ready cohort and restricting to participants with a valid PREVENT risk estimate, the analytic sample comprised 30,134 participants (1,680 events, 28,454 non-events). Variables were harmonized to match PREVENT inputs, including re-coding sex and converting binary predictors (blood-pressure treatment, statin use, diabetes, and smoking) to numeric indicators. IMM-AGE was analyzed as its raw pre-computed z-score. Ten-year cardiovascular risk was estimated using the PREVENT model as implemented in the preventr package (v0.11.0, Mayer et al; 2025), based on age, sex, systolic blood pressure, antihypertensive treatment, total cholesterol, HDL cholesterol, statin use, diabetes, smoking, estimated glomerular filtration rate, and body mass index. Predicted risks were converted to log-odds using log(risk / (1 - risk)) to serve as an offset in subsequent models. Participants with missing PREVENT risk estimates were excluded.

To evaluate the incremental contribution of IMM-AGE, we performed a sex-stratified bootstrap offset analysis. Within each sex stratum (n total female = 16622, cases female = 671, n total male = 13512, cases male = 1009), we performed case-anchored control-resampling bootstrap in which all cases were retained and controls were sampled without replacement at a 4:1 control-to-case ratio. In each bootstrap iteration, a logistic regression model was fitted using the glm function, with event status as the outcome, the PREVENT log-odds included as an offset term, and IMM-AGE as the sole predictor without an intercept. This procedure was repeated for 1,000 iterations, and only converged models were retained for downstream summaries.

Net reclassification improvement^49^ (NRI) was calculated within each bootstrap iteration by comparing predicted risk from the base PREVENT model with that from the PREVENT-plus-IMM-AGE model. Risk categories were defined using standard PREVENT thresholds: low (<3%), borderline (3-5%), intermediate (5-10%), and high (≥10%). Event NRI was defined as the proportion of cases moving to a higher minus lower risk category, and non-event NRI as the proportion of controls moving to a lower minus higher category; total NRI was defined as their sum.Point estimates of sex-specific NRI were computed from the mean predicted risk across bootstrap iterations, applied to the standard reclassification categories. Associated 95% intervals and two-sided p values were derived from the corresponding bootstrap distributions.

#### Risk score change by baseline PREVENT category - Events

Among individuals who experienced a cardiovascular event within 10 years, the magnitude and direction of risk score change when adding IMM-AGE to PREVENT were examined across four baseline risk categories: Low (<3%), Borderline (3–5%), Intermediate (5–10%), and High (≥10%). The augmented risk score was computed as plogis(PREVENT logit + β × IMM-AGE Z-score), where β is the sex-specific bootstrap median coefficient. This figure illustrates whether IMM-AGE preferentially upwardly reclassifies events who were initially underestimated by PREVENT.

#### Risk score change by baseline PREVENT category - Non-Events

The same risk change analysis was performed for individuals who did not experience a cardiovascular event. Among non-events, a beneficial reclassification is a downward shift in predicted risk. This figure examines whether IMM-AGE appropriately lowers estimated risk for non-events across baseline PREVENT risk strata, and whether any categories show net harmful upward reclassification.

#### Reclassification sensitivity and specificity across risk thresholds

Using the standard PREVENT risk categories (Low <3%, Borderline 3–5%, Intermediate 5–10%, High ≥10%) as decision thresholds, sensitivity and specificity of upward reclassification were computed across bootstrap iterations. Sensitivity reflects the proportion of true events whose risk category was increased or maintained (i.e., not reclassified downward) when IMM-AGE was added; specificity reflects the proportion of non-events whose category was decreased or maintained (i.e., not reclassified upward). Results are presented with bootstrap confidence intervals stratified by sex.

#### Lipid-Lowering Treatment reclassification: sensitivity and specificity

A clinically oriented reclassification analysis was performed using the lipid-lowering treatment (LLT) eligibility threshold - a 10-year risk of at least 5% (Intermediate category or above) - as the decision boundary. Sensitivity among true events was computed under two definitions, both stratified by sex with bootstrap confidence intervals. The strict definition counted an event as correctly handled only if its risk category was increased or maintained, as in the threshold-based analysis above. The clinically adjusted definition additionally did not penalize downward reclassification that left an event at or above the LLT-eligibility threshold - such individuals remain treatment-eligible - so that only events reclassified below the threshold were counted as missed.

### TAVR cohort

#### Projection of TAVR samples onto the reference manifold

Flow cytometry data from TAVR patients were prepared using the same set of immune features as described above, and standardized in an identical manner. These samples were projected onto the learned diffusion space using the dm_predict() function from the destiny package (v 3.18, Angerer et al; 2016), with the distance metric set to Minkowski (p = 1.5).

Projected coordinates (DC1 and DC2) were merged with the original embedding to visualize the relative positioning of TAVR samples in relation to the Ellison-derived manifold.

#### Estimation of IMM-AGE scores for projected samples

To assign IMM-AGE scores to TAVR samples, we computed each sample’s three nearest neighbors (k = 3) among the reference Ellison samples using the nn2() function from the RANN package (v2.6.2), based on DC1/DC2 coordinates. Each TAVR sample’s IMM-AGE score was estimated as the mean of its nearest neighbors’ DPT values. This pseudotime-based projection approach yielded continuous IMM-AGE estimates that were directly comparable across cohorts.

#### Validation of projection-based IMM-AGE estimation

To assess the fidelity of the projection method, we evaluated its ability to recapture known IMM-AGE scores for Ellison samples with independently profiled values. Using ggplot2, we visualized diffusion map coordinates by sample origin - Elison Trajectory, TAVR samples, HIMC controls, and selected reanalyzed healthy individuals from the SELA cohort (**Supplementary Fig. 5C**). We also visualized the distributions of IMM-AGE scores, colored by the origin of the samples to show comparable distributions of the scores (**Supplementary Fig. 5D**). Finally, we plotted the projected scores (IMM-AGE Projection) against the reference scores from the published Ellison dataset (IMM-AGE Original) for matched individuals. The resulting correlation coefficient (ρ = 0.878) confirmed strong agreement between the two sets of values, supporting the validity of the IMM-AGE score transfer via nearest-neighbour matching (**Supplementary Fig. 5E**).

#### IMM-AGE score adjustment for clinical confounders

To adjust the IMM-AGE scores of TAVR study participants for potential clinical and demographic confounders, we used ridge regression (L2-penalized) implemented via cv.glmnet() from the glmnet R package. The model included the following covariates: age, sex, body mass index (BMI), NT-proBNP, estimated glomerular filtration rate (CKD stage ≥3), atrial fibrillation (AF), neutrophil-to-lymphocyte ratio, NYHA class, STS score, metabolic syndrome, and cardiovascular disease history. The regularization parameter (lambda.min) was selected via 10-fold cross-validation. The adjusted IMM-AGE score was defined as the residual from this model (i.e., the difference between observed and predicted IMM-AGE), capturing variation not explained by the covariates.

#### Clustering of baseline and 1-Month post-procedure echocardiographic profiles

To identify subgroups of patients with distinct baseline and 1-month post-procedure cardiac phenotypes, we performed unsupervised clustering based on four pre-procedure echocardiographic parameters: global longitudinal strain (GLS), biplane left ventricular ejection fraction (LVEF), left ventricular mass index (LVMI), and right ventricular systolic pressure (RVSP). Only individuals with complete data across these features were included.

Principal component analysis (PCA) was conducted using the prcomp() function in base R, with variables scaled before decomposition. The first two principal components, explaining the majority of variance, were used as input for clustering. The optimal number of clusters was estimated via the Optimal_Clusters_KMeans() function from the ClusterR package, using the distortion-fK criterion.

K-means clustering was subsequently performed using eclust() from the factoextra package, specifying k= 2 clusters. Cluster assignment and separation were visualized in PCA space using ggplot2. Cluster-specific distributions of echocardiographic parameters were compared using boxplots and independent t-tests (stat_compare_means() from ggpubr) (**Supplementary Fig. 5A-B, Supplementary Tables 14 and 15**).

#### Nonparametric testing of cluster-based risk

To assess whether echocardiographic cluster assignment at baseline or at 1 month was associated with subsequent patient mortality, we performed non-parametric permutation testing in R, using the dplyr package (v1.1.4, Wickham et al. 2026) for data preprocessing. For each time point, we restricted analyses to individuals belonging to the two predefined echocardiographic clusters and recorded mortality as a binary variable. We computed the observed difference in mortality proportions between the two groups and generated a null distribution by permuting cluster labels 500 times, recalculating the difference at each iteration. Two-sided permutation p-values were obtained as the proportion of permuted statistics whose absolute value exceeded the observed effect. This procedure provides a distribution-free evaluation of whether mortality risk differs between echocardiographic phenotypes.

#### Logistic regression modelling of 1-month echocardiographic phenotype

To evaluate whether baseline IMM-AGE independently predicted maladaptive cardiac response following TAVR, we constructed a multivariable logistic regression model using the 1-month echocardiographic phenotype as the binary outcome. Candidate predictors included IMM-AGE, demographic factors (age, sex), clinical variables (BMI, metabolic syndrome, cardiovascular comorbidity, atrial fibrillation, CKD stage ≥3, NYHA class, STS score), and inflammatory or cardiac biomarkers (NLR, NT-proBNP). Continuous variables were standardized prior to modelling, and categorical variables were encoded as fixed-effect factors. Model coefficients were visualized using standardized effect plots, generated with the sjPlot package (v2.9.0).

To assess whether IMM-AGE provides predictive information beyond baseline cardiac structure and function, we fitted a multivariable logistic regression model for one-month echocardiographic outcome. The model included IMM-AGE, baseline echocardiographic phenotype (favorable vs. unfavorable), and the same demographic, clinical, and laboratory covariates used in prior analyses. Continuous variables were standardized prior to modeling, and categorical variables were encoded as factors. To mitigate small-sample and separation bias, regression coefficients and inference were obtained using Firth’s penalized likelihood approach implemented in the logistf R package (v. 1.26.1). Initial model specification and multicollinearity diagnostics were performed using base R (stats) and the car package (v 3.1-3). Data manipulation was conducted using dplyr (v.1.1.4), and model visualization was generated using ggplot2 (v.4.0.1). Model coefficients are reported as log-odds with corresponding confidence intervals.

#### Laboratory and clinical Discrimination of Post-TAVR Adaptation that

To further investigate predictors of post-TAVR adaptation, we analyzed a panel of clinical and molecular variables grouped into three domains: laboratory biomarkers (Gal-3, GDF15, NT-proBNP, CRP, hs-TnI), clinical predictors (Age, NYHA class, STS score), and echocardiographic severity indices (AVA and Mean Aortic Gradient). Relevant values were extracted and merged using unique biobanking identifiers. All variables were compared between 1-month adaptation phenotypes within the subgroup of patients with an unfavorable echocardiographic profile at baseline. Group differences were assessed using the t test (ggpubr::stat_compare_means, v0.6.0) and visualized with ggplot2 (v.4.0.1) and patchwork (v1.2.0) for composite figure construction.

### PBHS cohort

#### Selection of PBHS patients and preprocessing of immune features for IMM-AGE Estimation

To characterize immune aging, we selected six cytometry-derived immune cell populations previously shown to be informative for IMM-AGE dynamics: effector memory CD4⁺ T cells, naïve CD4⁺ T cells, effector CD8⁺ T cells, naïve CD8⁺ T cells, and effector memory CD8⁺ T cells. Although the original IMM-AGE panel included seven populations, two were unavailable in our PBHS panel. Therefore, we simulated the reduced panel on the original SELA cohort to evaluate its sufficiency (**Supplementary Fig. 6F**). To mitigate the impact of outliers, each immune feature was scaled using a robust z-score computed over the central 80% (10th-90th percentile) of its distribution.

#### Construction of the reference immune aging trajectory

To uncover the intrinsic structure of immune variation, we applied diffusion maps using the DiffusionMap() function from the destiny package (v3.18, Angerer et al; 2016). The scaled cytometry data matrix was embedded into a low-dimensional manifold. Subsequently, diffusion pseudotime (DPT) scores were derived using *DPT* function to reflect the continuum of immune aging. These DPT values served as IMM-AGE scores for the reference (Ellison) cohort.

#### Permutation-based assessment of projection stability

To ensure robust IMM-AGE estimation and assess sampling variance, we implemented a 100-iteration permutation framework. In each iteration, PBHS samples were randomly partitioned into subsets (∼100 samples per batch). For each subset, IMM-AGE scores were re-estimated by projecting onto the same reference manifold and applying the same nearest-neighbor assignment. The distribution of IMM-AGE values across permutations was summarized by computing the median and standard deviation per individual.

#### Projection of PBHS Samples onto the Reference Manifold

Flow cytometry data from PBHS individuals were processed using the same set of immune features and normalization strategy. Using dm_predict() from the destiny package (distance metric: Minkowski, p = 1.5), these samples were projected onto the learned diffusion space. Visualization of projected and reference samples was performed using the ggplot2 R package (v4.0.1; Wickham et al., 2016).

#### Estimation of IMM-AGE scores for projected samples

For each projected sample, we computed its three nearest neighbors (k = 3) among the reference Ellison samples in DC1/DC2 space using the nn2() function from the RANN package (version 2.6.2). IMM-AGE scores were then estimated as the average DPT score of these neighbors. This pseudotime-projection approach enabled a continuous and cohort-comparable IMM-AGE quantification.

#### Physiological and clinical risk correlates of IMM-AGE

IMM-AGE was compared across ordered categories of coronary artery calcium and diastolic function. Differences between adjacent categories were assessed using two-sided t-tests implemented with *stat_compare_means* from *ggpubr* (v0.6.0; Kassambara et al., 2026). Data wrangling and visualization were performed using *dplyr* (v1.1.4; Wickham et al., 2026) and *ggplot2* (v.4.0.1; Wickham et al., 2016).

#### Cardiovascular events in PBHS

In the project Baseline Health Study, cardiovascular events were collected using clinical case report forms during scheduled annual or study visits. Outcome curation was completed on November 20th 2023. A total of 129 cardiovascular events were recorded over a median follow-up of 1463 days. First events consisted of atherosclerotic events (n= 67), incident atrial fibrillation or atrial flutter (n=33) heart failure or hypertensive emergency (n=10), thrombotic events (n=10), cardiovascular intervention (n=8) and cardiovascular death (n=1). 29 patients died without experiencing cardiovascular events, with the exception of one patient with cancer, the cause of the other 28 events was not clearly identified at the time of outcome curation.

#### Similarity-based matching and modeling

To control for clinical confounding, we constructed a similarity network among participants using Gower distance computed across 15 demographic, clinical, laboratory, and treatment covariates: age at enrollment, sex, HDL cholesterol, LDL cholesterol, C-reactive protein (CRP), neutrophil-to-lymphocyte ratio (NLR), systolic blood pressure, glomerular filtration rate (GFR), body mass index (BMI), history of cardiovascular disease, diabetes, cerebrovascular disease, lipid-lowering medication use, smoking status, and antihypertensive medication use. Binary variables (sex, cardiovascular history, diabetes, cerebrovascular disease, lipid-lowering medication use, and antihypertensive medication use) were treated as asymmetric binary features in the Gower distance calculation, such that shared presence of a condition was weighted more heavily than shared absence.

Each case was matched to its 10 most similar controls by minimum Gower distance, yielding an initial set of case-centered clusters. Clusters sharing five or more controls were subsequently merged using a graph-based connected-component approach, in which clusters were represented as nodes and edges were drawn between pairs sharing sufficient controls. This merging step unified overlapping neighborhoods into larger matched case-control clusters. Controls appearing in multiple clusters following merging were deduplicated, retaining a single membership per merged cluster. A total of 54 merged clusters were retained for downstream analysis. Participants with missing values in any of the 15 matching covariates were excluded prior to Gower distance computation; of 122 individuals who experienced a cardiovascular event, 105 cases and 844 controls had complete covariate data and were retained for analysis.

Conditional logistic regression stratified by merged cluster ID was used as the primary model for the matched case-control analysis, as this is the canonical approach for matched case-control designs and directly conditions on cluster membership without requiring estimation of a random effect variance. The model included age, sex, HDL cholesterol, LDL cholesterol, systolic blood pressure, CRP, NLR, eGFR, BMI, history of cardiovascular disease, cerebrovascular disease, diabetes, smoking status, and antihypertensive medication use as fixed effects. Lipid-lowering medication use was excluded from regression models due to near-complete separation in the matched sample (1 of 949 matched individuals had lipid medication use), rendering stable coefficient estimation impossible. Model fit was compared between models with and without IMM-AGE using a likelihood ratio test.

#### Assessment of case-control similarity within clusters

To evaluate matching quality, we computed pairwise Gower distances between all case-control pairs using the same 15 covariates that entered the matching procedure, and converted distances to similarity scores (similarity = 1 − distance). Case-control pairs were categorized as either within-cluster (case and control belonging to the same merged cluster) or between-cluster (case and control from different clusters). By focusing exclusively on case-control comparisons, we ensured that similarity was assessed in the context most relevant to the outcome analysis. We compared the distributions of within-and between-cluster similarity scores using a two-sample t-test to determine whether cases were more similar to their matched controls than to unmatched controls.

#### Assessment of covariate balance within clusters

To assess covariate balance between cases and their matched controls, we computed cluster-aware standardized mean differences (SMDs) for all 15 covariates. For each variable, the within-cluster SMD was computed as the difference between the cluster-level case mean and control mean, divided by the global pooled standard deviation across all individuals. This denominator was computed globally rather than within-cluster to avoid instability in clusters with few individuals. The per-cluster SMDs were then averaged across clusters, weighted by the harmonic mean of the number of cases and controls within each cluster, which downweights clusters with highly unequal case-to-control ratios. Results were visualized as a Love plot with a conventional balance threshold of SMD = 0.1.

For continuous variables — including age at enrollment, HDL cholesterol, LDL cholesterol, CRP, NLR, systolic blood pressure, eGFR, and BMI - we additionally computed the mean value of each variable among controls in each cluster and compared it to the corresponding case mean using paired t-tests aggregated across clusters. For binary variables — including sex, history of cardiovascular disease, diabetes, cerebrovascular disease, lipid-lowering medication use, smoking status, and antihypertensive medication use — we identified the modal value among controls within each cluster and computed the proportion of cases whose value matched that mode (agreement rate).

#### Comparison of IMM-AGE between cases and controls

To evaluate differences in IMM-AGE scores between cases and controls, we extracted all case-control pairs from the matched dataset and retained only individuals with non-missing IMM-AGE values. For each matched cluster, we computed the mean IMM-AGE separately for cases and controls. The overall difference between cases and controls was then calculated by averaging the per-cluster differences. To assess statistical significance, we performed a permutation test by randomly flipping the case/control labels within each cluster (i.e., role-swapping) across 1,000 iterations. Permutation p-values were computed as the proportion of permuted differences whose absolute value exceeded the observed difference.

#### Comparison of IMM-AGE differences: case-control vs. control-control pairs

To test whether IMM-AGE differences between cases and controls exceeded those observed between matched controls, we conducted a paired difference analysis at the cluster level. Within each cluster, all case-control IMM-AGE differences were computed and averaged, as were all pairwise IMM-AGE differences between controls (control-control). Clusters with at least one case and at least two controls were included. The per-cluster difference between the two types of comparisons (case-control minus control-control) was then averaged across clusters. A paired permutation test was used to assess statistical significance by randomly flipping the sign of these differences within clusters across 1,000 iterations. The p-value was calculated as the fraction of permutations with a mean difference greater than or equal to the observed value.

#### COX proportional hazards modeling

Associations between IMM-AGE and incident cardiovascular events were evaluated using Cox proportional hazards regression. Time-to-event was defined from baseline to the first occurrence of major adverse cardiovascular events (MACE). Models were adjusted for established cardiovascular risk factors, including age, sex, LDL cholesterol, systolic blood pressure, C-reactive protein, neutrophil-to-lymphocyte ratio, estimated glomerular filtration rate, body-mass index, prior cardiovascular disease, diabetes, smoking status, and antihypertensive medication use. To account for the matched structure of the dataset, a random intercept for matched clusters (merged_cluster_id) was included using a mixed-effects Cox model implemented with the *coxme* function from the *coxme* R package (v2.2-22, Therneau et al. 2024^44^).

#### Standardization and model output visualization

Continuous covariates were standardized (mean = 0, standard deviation = 1) prior to model fitting to facilitate comparison of relative effect sizes across the conditional logistic regression model. Odds ratios (ORs) with 95% confidence intervals were derived from the standardized model coefficients and visualized using the sjPlot R package (version 2.9.0).

#### Model comparison via likelihood ratio test

To assess whether inclusion of IMM-AGE significantly improved model fit, we compared two nested conditional logistic regression models using a likelihood ratio test implemented via the anova() function with test = “Chisq”. The full model included IMM-AGE alongside all clinical and demographic covariates; the reduced model was identical but excluded IMM-AGE. Both models were fitted using the clogit function from the survival R package, stratified by merged cluster ID. The difference in log-likelihoods between the two models was evaluated as a chi-squared statistic with one degree of freedom.

### Data visualization and figure assembly

Plots were generated in R v4.3.3 using ggplot2 and exported as vector graphics. Final figures were assembled, arranged, and annotated in Adobe Illustrator 2025 (Adobe Inc., San Jose, CA,); no adjustments were made that altered the underlying data.

## Data availability

All publicly available gene-expression datasets analysed in this study were obtained from the Gene Expression Omnibus (GEO) under the following accession codes: GSE20681, GSE20680, GSE90074, GSE180081, GSE59867, GSE58294, GSE218474, GSE209567, GSE38267 and GSE33463, as noted on Figure 1F. Data from the SELA cohort will be shared by the corresponding author upon reasonable request. Gene-expression and methylation data from the FHS are available through dbGaP under study accession phs000007; the corresponding phenotypic data are available under accession codes pht003099 (gender and age at exam 8), pht000747 (smoking status, blood pressure and blood-pressure treatment), pht000742 (Total cholesterol and fasting glucose) and pht000041 (diabetes treatment), while cardiovascular-disease status at exam 8 and all-cause mortality during follow-up were derived from the files ‘survival and follow-up status for cardiovascular events’ (pht003316) and ‘survival – all-cause mortality’ (pht003317), respectively. UKB data are available through the UK Biobank repository, to approved researchers upon application to the UK Biobank access team; the analyses reported here were conducted under Application ID 1164153. Data from the TAVR cohort are provided in Supplementary Table 12. Data from the PBHS are available upon request to the Project Baseline Health Study team.

## Supplementary Note 1

### Robustness of the IMM-AGE signal after comprehensive clinical adjustment

To evaluate whether the association between IMM-AGE and post-procedural cardiac phenotypes is independent of established clinical risk markers, we performed an extended adjustment analysis incorporating a broad panel of pre-TAVR demographic, clinical, echocardiographic, and circulating biomarker variables.

All analyses were restricted to participants with complete data across the evaluated covariates to ensure consistency of adjustment. IMM-AGE was adjusted for age, functional status (NYHA class), surgical risk (STS score), baseline echocardiographic severity (aortic valve area and mean aortic gradient), and established biomarkers of myocardial stress, inflammation, and fibrosis, including NT-proBNP, high-sensitivity troponin I, CRP, Galectin-3, and GDF-15. This approach was designed to isolate the component of immune aging not attributable to conventional cardiovascular risk factors or disease severity.

Following adjustment, the residualized IMM-AGE signal remained significantly associated with the one-month echocardiographic phenotype, with higher adjusted IMM-AGE observed among individuals exhibiting maladaptive ventricular response at one month.

To further contextualize this finding within a multivariable clinical framework, we additionally fit a standardized logistic regression model including IMM-AGE and all aforementioned covariates. Effect sizes were visualized in a supplementary forest plot using standardized predictors to enable direct comparison across variables. Consistent with the residualization analysis, IMM-AGE remained an independent contributor to the prediction of adverse one-month echocardiographic phenotype after accounting for demographic, clinical, echocardiographic, and biomarker measures.

Collectively, these sensitivity analyses demonstrate that the association between IMM-AGE and early post-TAVR cardiac phenotype is not driven by traditional markers of disease severity, inflammation, or myocardial injury, supporting the interpretation of IMM-AGE as a distinct and clinically relevant immune-aging signal.

**Extended Data Figure 1:**
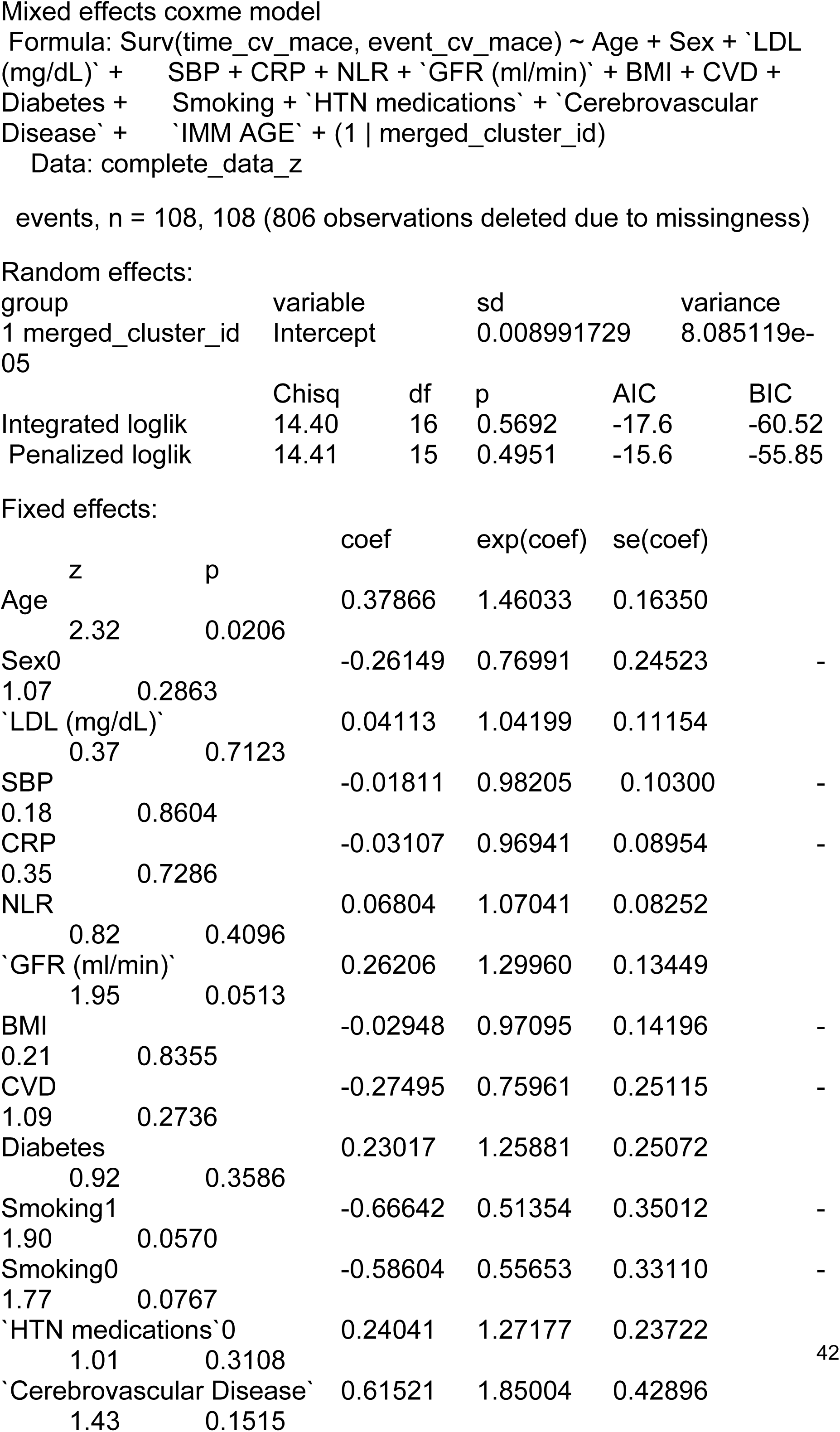
PBHS COX modeling

## References

1. Roth, G. A. et al. Global Burden of Cardiovascular Diseases and Risk Factors, 1990–2019: Update From the GBD 2019 Study. J. Am. Coll. Cardiol. 76, 2982–3021 (2020).

2. Libby, P. The changing landscape of atherosclerosis. Nature 592, 524–533 (2021).

3. D’Agostino, R. B. et al. General Cardiovascular Risk Profile for Use in Primary Care. Circulation 117, 743–753 (2008).

4. Goff, D. C. et al. 2013 ACC/AHA Guideline on the Assessment of Cardiovascular Risk. Circulation 129, S49–S73 (2014).

5. Khan, S. S. et al. Novel Prediction Equations for Absolute Risk Assessment of Total Cardiovascular Disease Incorporating Cardiovascular-Kidney-Metabolic Health: A Scientific Statement From the American Heart Association. Circulation 148, 1982–2004 (2023).

6. Arnett, D. K. et al. 2019 ACC/AHA Guideline on the Primary Prevention of Cardiovascular Disease: A Report of the American College of Cardiology/American Heart Association Task Force on Clinical Practice Guidelines. Circulation 140, e596–e646 (2019).

7. Libby, P. Inflammation in atherosclerosis. Nature 420, 868–874 (2002).

8. Hansson, G. K. & Hermansson, A. The immune system in atherosclerosis. Nat. Immunol. 12, 204–212 (2011).

9. Libby, P. & Hansson, G. K. From Focal Lipid Storage to Systemic Inflammation. JACC 74, 1594–1607 (2019).

10. Engelen, S. E., Robinson, A. J. B., Zurke, Y.-X. & Monaco, C. Therapeutic strategies targeting inflammation and immunity in atherosclerosis: how to proceed? Nat. Rev. Cardiol. 19, 522–542 (2022).

11. Liu, J. et al. Advances in pharmacological research on myocardial remodeling agents: A decade in review. Medicine (Baltimore) 104, e42757 (2025).

12. Davis, M. M. & Brodin, P. Rebooting Human Immunology. Annu. Rev. Immunol. 36, 843–864 (2018).

13. Kaczorowski, K. J., et al. Continuous immunotypes describe human immune variation and predict diverse responses. Proc. Natl. Acad. Sci. 114, E6097–E6106 (2017).

14. Sayed, N. et al. An inflammatory aging clock (iAge) based on deep learning tracks multimorbidity, immunosenescence, frailty and cardiovascular aging. Nat. Aging 1, 598–615 (2021).

15. Sparks, R. et al. A unified metric of human immune health. Nat. Med. 30, 2461–2472 (2024).

16. Fuentealba, M. et al. A blood-based epigenetic clock for intrinsic capacity predicts mortality and is associated with clinical, immunological and lifestyle factors. Nat. Aging 5, 1207–1216 (2025).

17. Climente-González, H. et al. Interpretable machine learning leverages proteomics to improve cardiovascular disease risk prediction and biomarker identification. Commun. Med. 5, 170 (2025).

18. Ho, F. K. et al. A Proteomics-Based Approach for Prediction of Different Cardiovascular Diseases and Dementia. Circulation 151, 277–287 (2025).

19. Alpert, A. et al. A clinically meaningful metric of immune age derived from high-dimensional longitudinal monitoring. Nat. Med. 25, 487–495 (2019).

20. Cipriano, A. et al. Immune aging biomarkers for clinical trials. Nat. Med. 32, 2368–2382 (2026).

21. Wenzl, F. A. et al. Inflammation in Metabolic Cardiomyopathy. Front. Cardiovasc. Med. 8, (2021).

22. Frąk, W. et al. Pathophysiology of Cardiovascular Diseases: New Insights into Molecular Mechanisms of Atherosclerosis, Arterial Hypertension, and Coronary Artery Disease. Biomedicines 10, (2022).

23. Suffee, N., Le Goff, W. & Chen, J. Editorial: Cardiometabolic diseases and inflammatory responses. Front. Immunol. 15, (2024).

24. Qian, Q. et al. CVD Atlas: a multi-omics database of cardiovascular disease. Nucleic Acids Res. 53, D1348–D1355 (2025).

25. Sun, B. B. et al. Plasma proteomic associations with genetics and health in the UK Biobank. Nature 622, 329–338 (2023).

26. Savitz, S. T. et al. Predicting short-term outcomes after transcatheter aortic valve replacement for aortic stenosis. Am. Heart J. 256, 60–72 (2023).

27. Hoffmann*, J., et al. Inflammatory Signatures are Associated with Increased Mortality After Transfemoral Transcatheter Aortic Valve Implantation. ESC Heart Fail. 7, 2597–2610 (2020).

28. Kim, J. B. et al. Growth Differentiation Factor 15 is Associated with Lack of Ventricular Recovery and Mortality following Transcatheter Aortic Valve Replacement. Circ. Cardiovasc. Interv. 10, e005594 (2017).

29. Libby, P., Ridker, P. M. & Hansson, G. K. Inflammation in Atherosclerosis: From Pathophysiology to Practice. J. Am. Coll. Cardiol. 54, 2129–2138 (2009).

30. Ridker, P. M. From C-Reactive Protein to Interleukin-6 to Interleukin-1. Circ. Res. 118, 145–156 (2016).

31. Gisterå, A. & Hansson, G. K. The immunology of atherosclerosis. Nat. Rev. Nephrol. 13, 368–380 (2017).

32. Sayed, N. et al. An inflammatory aging clock (iAge) based on deep learning tracks multimorbidity, immunosenescence, frailty and cardiovascular aging. Nat. Aging 1, 598–615 (2021).

33. Sparks, R. et al. A unified metric of human immune health. Nat. Med. 30, 2461–2472 (2024).

34. Martín, P. & Sánchez-Madrid, F. T cells in cardiac health and disease. J. Clin. Invest. 135, (2025).

35. Saigusa, R., Winkels, H. & Ley, K. T cell subsets and functions in atherosclerosis. Nat. Rev. Cardiol. 17, 387–401 (2020).

36. Arges, K. et al. The Project Baseline Health Study: a step towards a broader mission to map human health. Npj Digit. Med. 3, 84 (2020).

37. Gautier, L., Cope, L., Bolstad, B. M. & Irizarry, R. A. affy—analysis of Affymetrix GeneChip data at the probe level. Bioinformatics 20, 307–315 (2004).

38. Davis, S. & Meltzer, P. GEOquery: a bridge between the Gene Expression Omnibus (GEO) and BioConductor. Bioinformatics 14, 1846–1847 (2007).

39. Hänzelmann, S., Castelo, R. & Guinney, J. GSVA: gene set variation analysis for microarray and RNA-Seq data. BMC Bioinformatics 14, 7 (2013).

40. Dewey, M. Metap: Meta-Analysis of Significance Values. (2025).

41. Hofner, B. & Hothorn, T. Stabs: Stability Selection with Error Control. (2026).

42. Wickham, H. Ggplot2: Elegant Graphics for Data Analysis. (Springer-Verlag New York, 2016).

43. Angerer, P. et al. destiny: diffusion maps for large-scale single-cell data in R. Bioinformatics 32, 1243 (2015).

44. Therneau, T. M. Coxme: Mixed Effects Cox Models. (2024).

45. Buuren, S. van & Groothuis-Oudshoorn, K. mice: Multivariate Imputation by Chained Equations in R. J. Stat. Softw. 45, 1–67 (2011).

46. Lindeman, R. H. Introduction to Bivariate and Multivariate Analysis. (Scott, Foresman, Glenview, Ill., 1980).

47. Groemping, U. Relative Importance for Linear Regression in R: The Package relaimpo. J. Stat. Softw. 17, 1–27 (2007).

48. Blanche, P., Dartigues, J.-F. & Jacqmin-Gadda, H. Estimating and comparing time-dependent areas under receiver operating characteristic curves for censored event times with competing risks. Stat. Med. 32, 5381–5397 (2013).

49. Kerr, K. F. et al. Net Reclassification Indices for Evaluating Risk Prediction Instruments: A Critical Review. Epidemiology 25, 114 (2014).

