## Supplementary Figure 6 for "Immune Aging is an Independent Risk Factor for Cardiovascular Disease" (from another version of this preprint: version 1, 2026-05-20)

**Supplementary Figure 6. Technical validation of network-based matching in PBHS cohort**

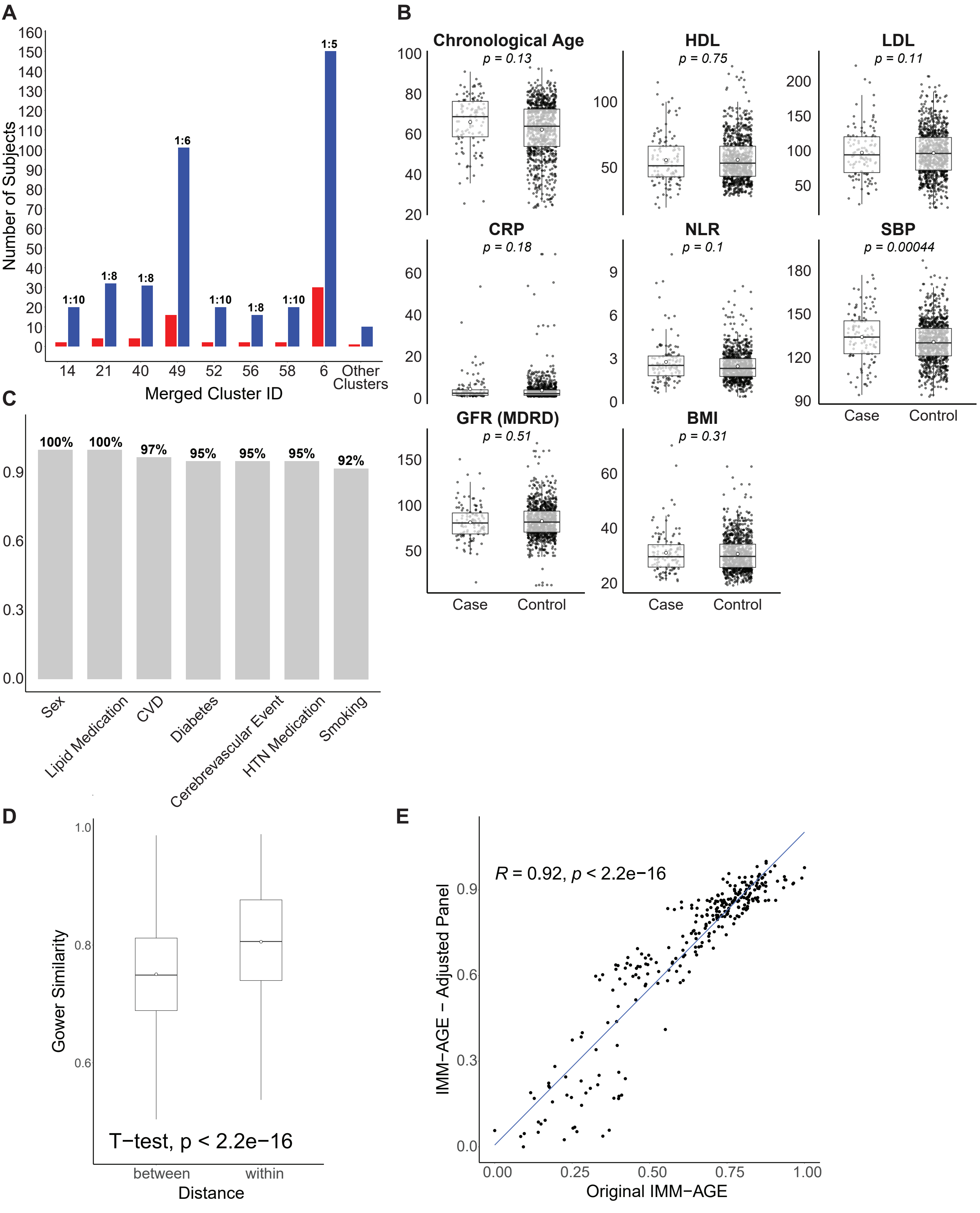

**Supplementary FIgure 6. Technical validation of network-based matching in PBHS cohort**
**(A)** Distribution of cardiovascular event cases and controls across matched clusters. Bar plot showing the composition of the 62 merged clusters generated by similarity-based matching. Each case was matched to its ten most clinically similar controls across 13 clinical and laboratory variables, and overlapping matched sets were merged into connected components. Blue bars indicate controls and red bars indicate cases. **(B)** Continuous covariate balance between cases and matched controls. Cluster-level comparison of continuous clinical and laboratory variables used for matching. Box plots show no substantial differences between cases and controls across matched clusters (t test). **(C)** Matching quality for binary clinical variables. Bar plot showing agreement between cases and the modal value of matched controls for categorical covariates, including sex, diabetes, smoking and medication use. Agreement rate was calculated as the proportion of cases whose value matched the modal value of their matched controls. **(D)** Intra- versus inter-cluster similarity. Box plot comparing pairwise Gower similarity for case-control pairs within the same matched cluster versus different clusters. Similarity was higher within clusters than between clusters (t test, $P < 2.2\times10^{-16}$). (E) Recapture of the original IMM-AGE signal using the PBHS-compatible immune panel. Correlation between original CyTOF-derived IMM-AGE scores and scores re-estimated using the five immune populations available in PBHS, validated in 46 SELA samples ($\rho = 0.92$, $P < 2.2\times10^{-16}$).
