## Supplementary Figure 5 for "Immune Aging is an Independent Risk Factor for Cardiovascular Disease" (from another version of this preprint: version 1, 2026-05-20)

### Supplementary Figure 5. Derivation of cardiac recovery phenotypes in the TAVR cohort

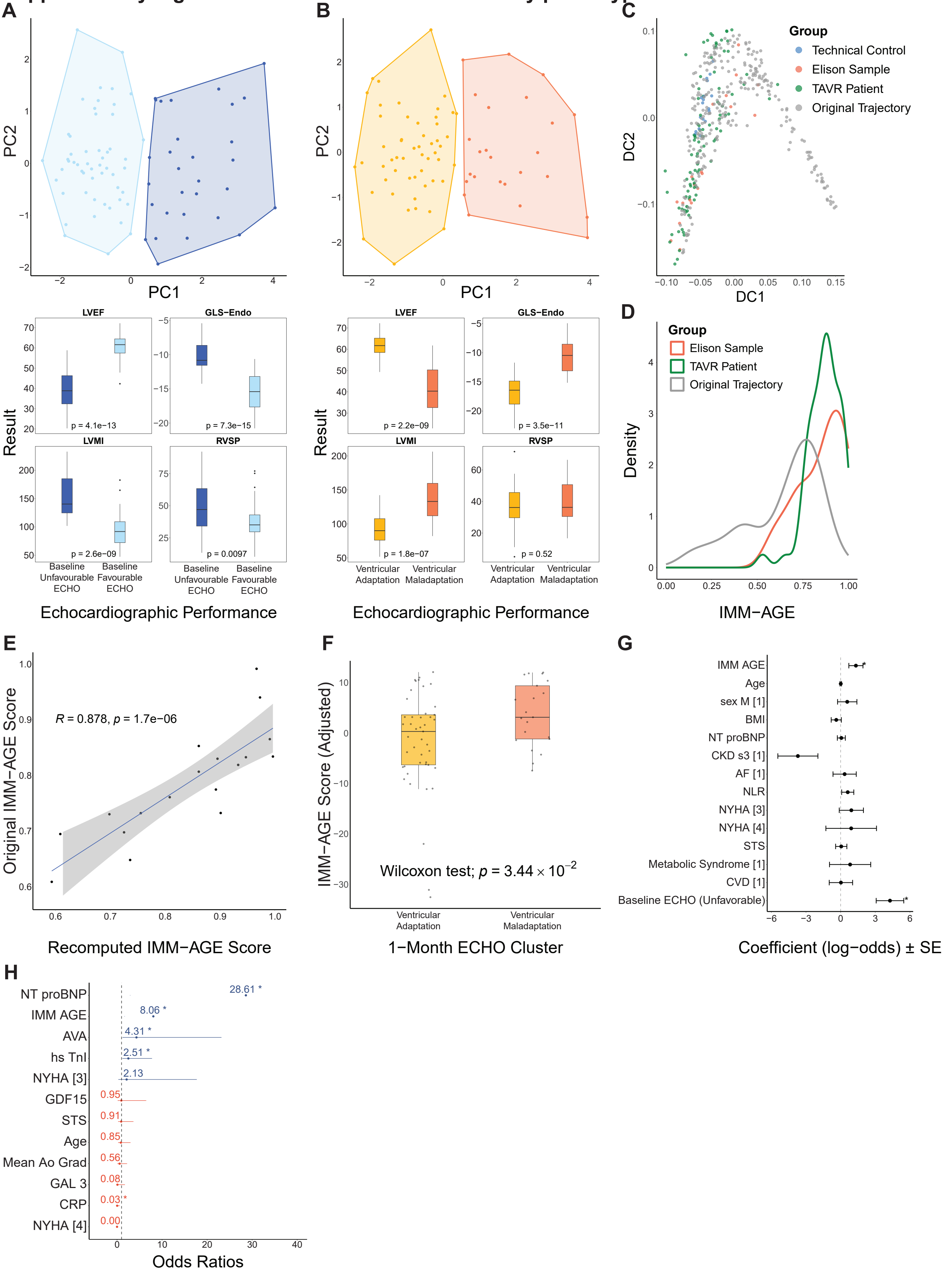

**Supplementary Figure 5. Derivation of cardiac recovery phenotypes in the TAVR cohort**

**(A,B)** Data-driven derivation of cardiac phenotypes. Principal component analysis of longitudinal echocardiographic parameters used to define ventricular adaptation and maladaptation states before and 1 month after TAVR. Box plots show selected echocardiographic features distinguishing the resulting groups. **(C)** Integration of clinical samples and internal controls into the immune-aging manifold. Diffusion map projection of the TAVR cohort onto the reference Ellison trajectory, together with re-analyzed SELA samples and technical controls. **(D)** Distribution of IMM-AGE scores in the reference and TAVR cohorts. Density plot comparing IMM-AGE score distributions in the Ellison reference cohort and the TAVR validation cohort. **(E)** Concordance between flow cytometry- and mass cytometry-derived IMM-AGE scores. Correlation between CyTOF-derived IMM-AGE and estimates obtained by flow cytometry in re-analyzed SELA samples ($\rho = 0.878$, $P < 2.2\times10^{-16}$). **(F)** Independence of IMM-AGE from conventional clinical and inflammatory risk factors. Boxplot comparing adjusted IMM-AGE scores (y-axis) across 1-month cardiac phenotypes (x-axis). The association remains significant (two-sided Wilcoxon test, $P = 0.034$) after regressing out the effects of age, STS score, NLR, and renal function. **(G)** Prognostic independence of IMM-AGE from baseline cardiac function. Forest plot of a logistic regression model identifying baseline IMM-AGE as a significant predictor of 1-month maladaptation (log-odds = 1.32, $p = 0.049$). The association remains significant after accounting for baseline echocardiographic performance (log-odds = 4.27, $p = 2.97\times10^{-5}$). **(H)** IMM-AGE association after multivariable adjustment. Forest plot of odds ratios from a model including IMM-AGE together demographic, cardiac and laboratory measurements. Baseline IMM-AGE is a significant predictor of 1-month maladaptation in this setting.
