## Supplementary Figure 4 for "Immune Aging is an Independent Risk Factor for Cardiovascular Disease" (from another version of this preprint: version 1, 2026-05-20)

### Supplementary Figure 4. Translation of IMM-AGE measurement from CyTOF to flow cytometry

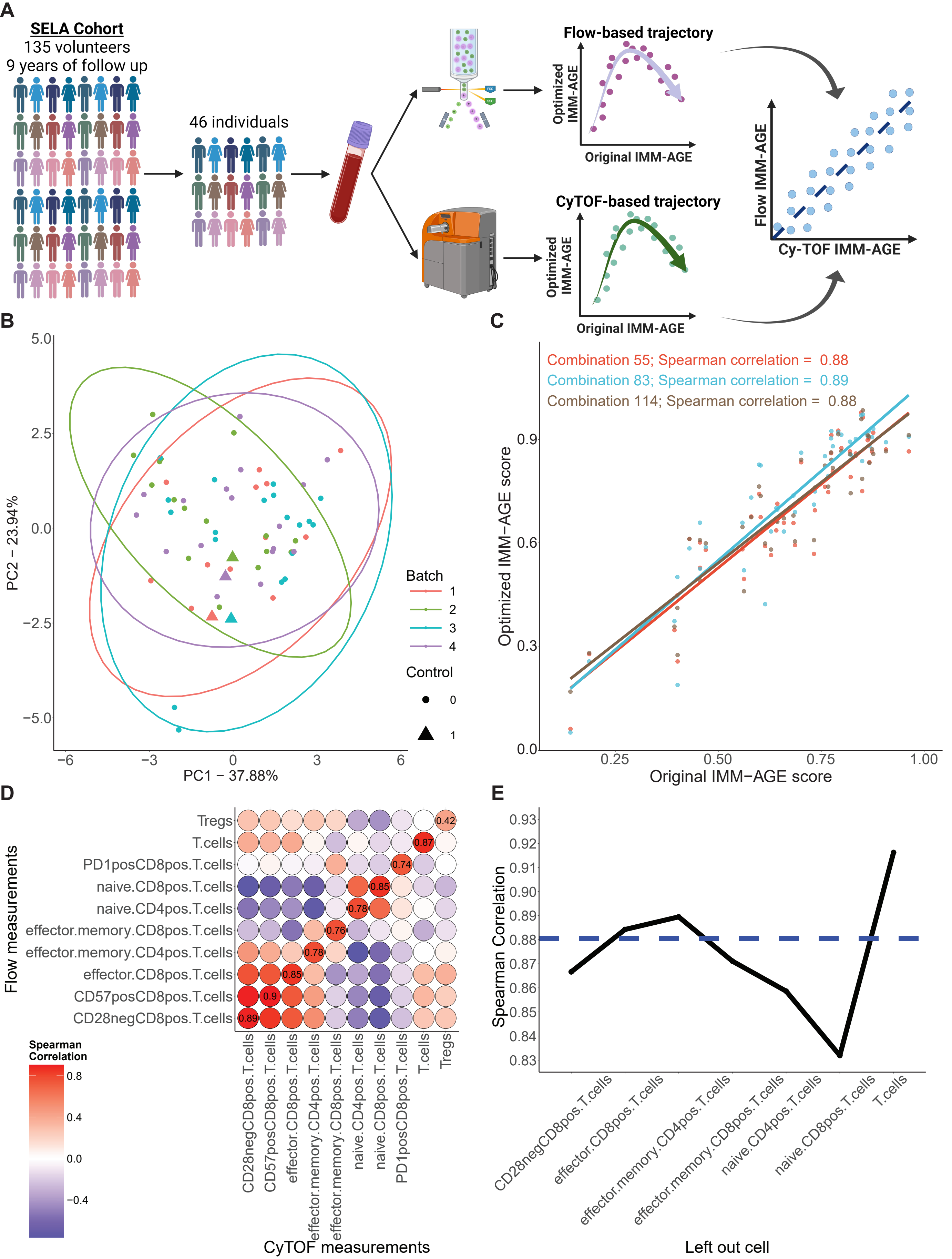

**Supplementary Figure 4. Translation of IMM-AGE measurement from CyTOF to flow cytometry.** **(A)** Cross-platform validation design. Schematic of the validation workflow in a subset of SELA participants (n = 46) re-profiled by standard flow cytometry. **(B)** Technical variation across flow cytometry batches. Principal component analysis of immune cell subsets across four processing batches. **(C)** Concordance of optimized panels across profiling platforms. Correlation of CyTOF- and flow cytometry-based IMM-AGE estimates for combinations 55, 83 and 114 in a subset of SELA participants (n = 46; $\rho = 0.88$, 0.89 and 0.88, respectively). **(D)** Concordance between flow cytometry and CyTOF measurements across selected immune populations. Flow and CyTOF measurements were highly correlated. **(E)** Robustness of IMM-AGE estimation to single-cell-type exclusion. Spearman correlation between original CyTOF-derived IMM-AGE scores and flow-based estimates after sequential exclusion of individual cell populations. High concordance was maintained across exclusions (minimum $\rho > 0.83$).
