## Supplementary Figure 3 for "Immune Aging is an Independent Risk Factor for Cardiovascular Disease" (from another version of this preprint: version 1, 2026-05-20)

### Supplementary Figure 3. In-silico validation of IMM-AGE optimized flow-cytometry panel

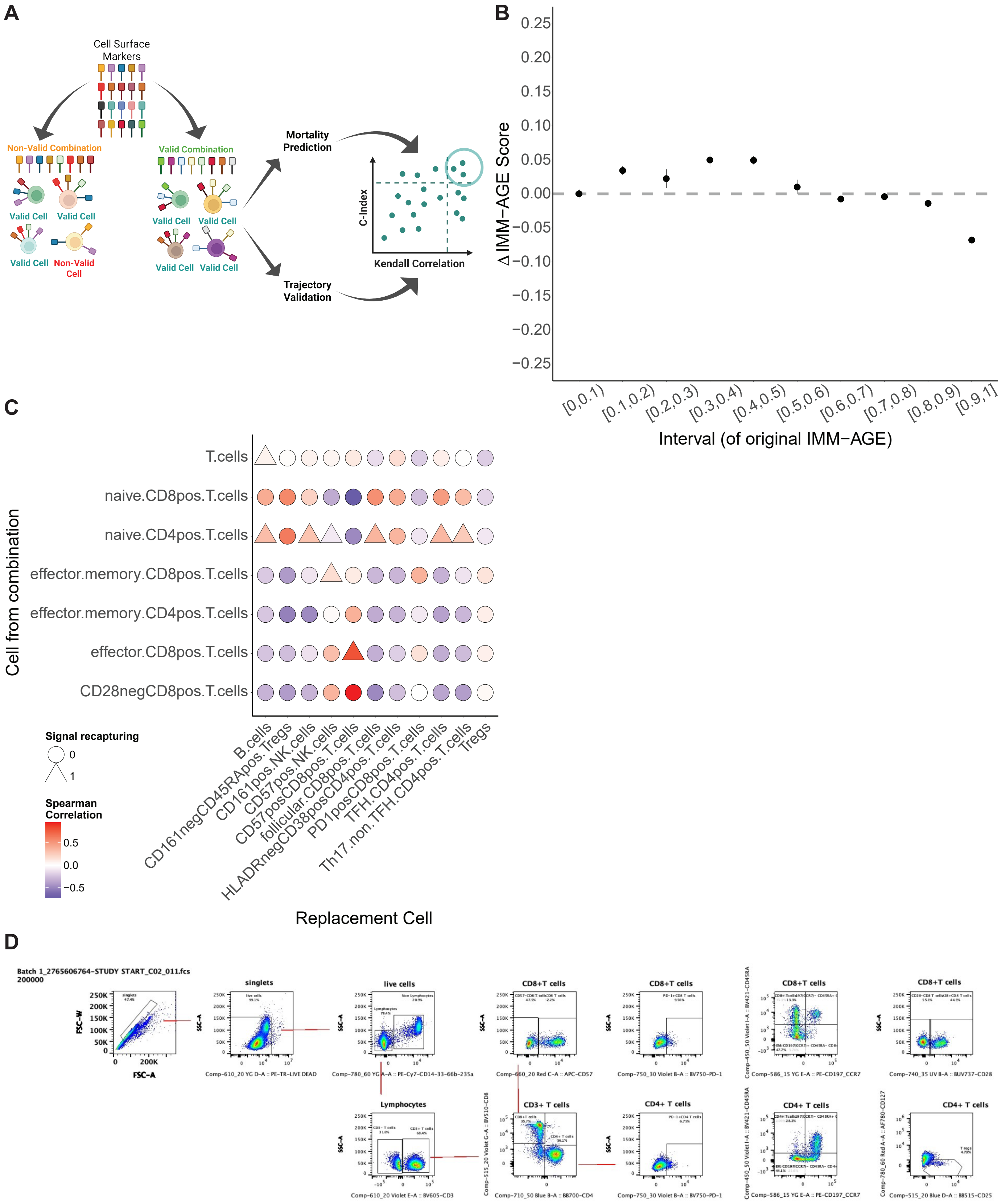

**Supplementary Figure 3. Computational selection and robustness of reduced-marker IMM-AGE panels. (A)** High-throughput simulation of reduced-marker panels. Schematic of the workflow used to evaluate all possible eight-marker combinations in the SELA CyTOF dataset by concordance with the original IMM-AGE trajectory and mortality prediction in the Framingham Heart Study. **(B)** Estimation accuracy across the immune-aging spectrum. Mean deviation (ΔIMM-AGE, y axis) between the original full-panel and optimized reduced-panel estimates across intervals of the original IMM-AGE score (x axis). Minimal deviation across intervals indicates stable performance throughout the IMM-AGE range. **(C)** Robustness of IMM-AGE signal to systematic subset replacement. Matrix showing the effect on signal recapture (Spearman correlation) of replacing individual cell types within the selected optimized combination with alternative immune subsets. Correlation values remained stable across replacements. (D) Flow cytometry gating strategy.
