## Supplementary Figure 2 for "Immune Aging is an Independent Risk Factor for Cardiovascular Disease" (from another version of this preprint: version 1, 2026-05-20)

### Figure 1. IMM-AGE is a quantifiable metric of cardiovascular morbidity and mortality risk.

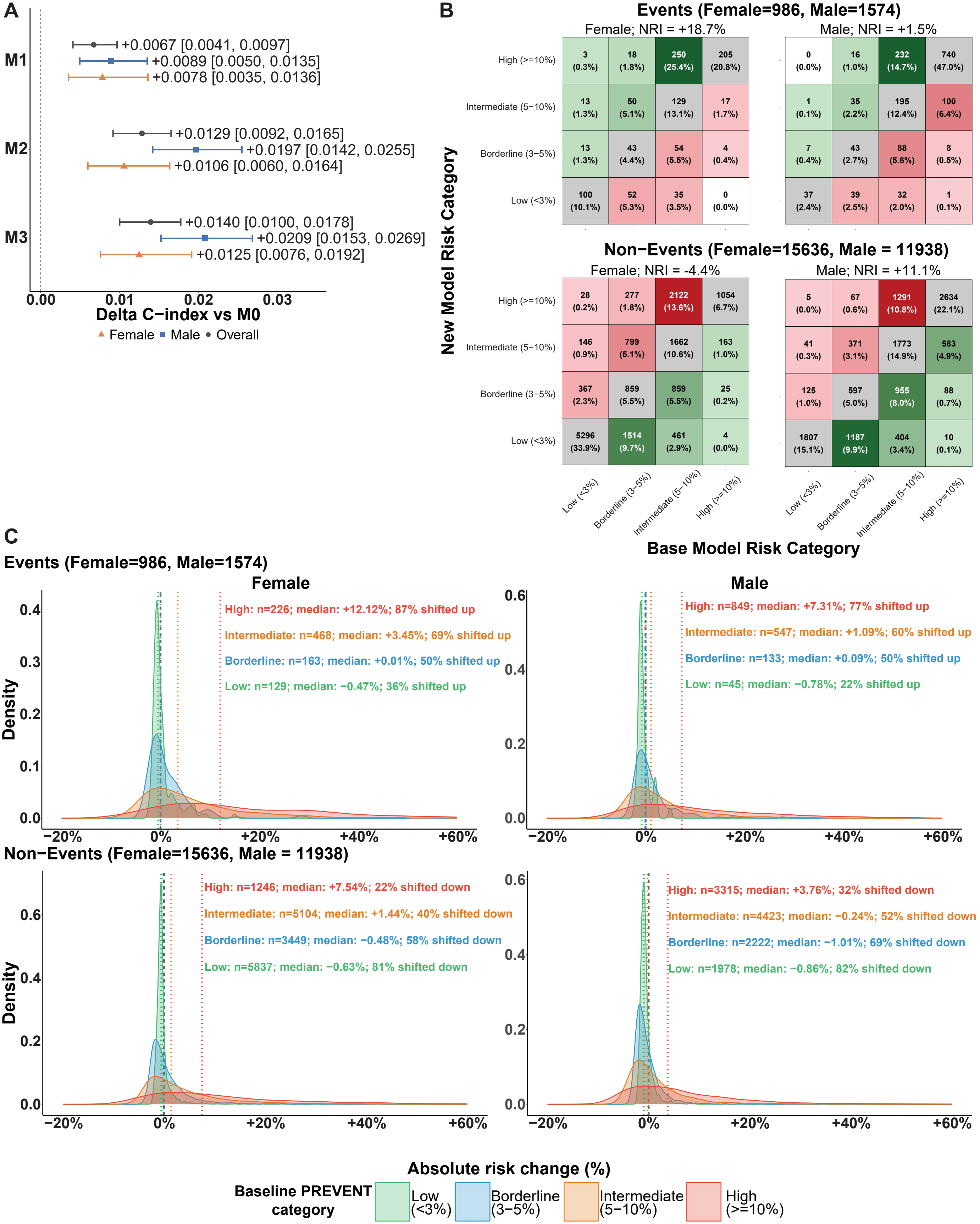

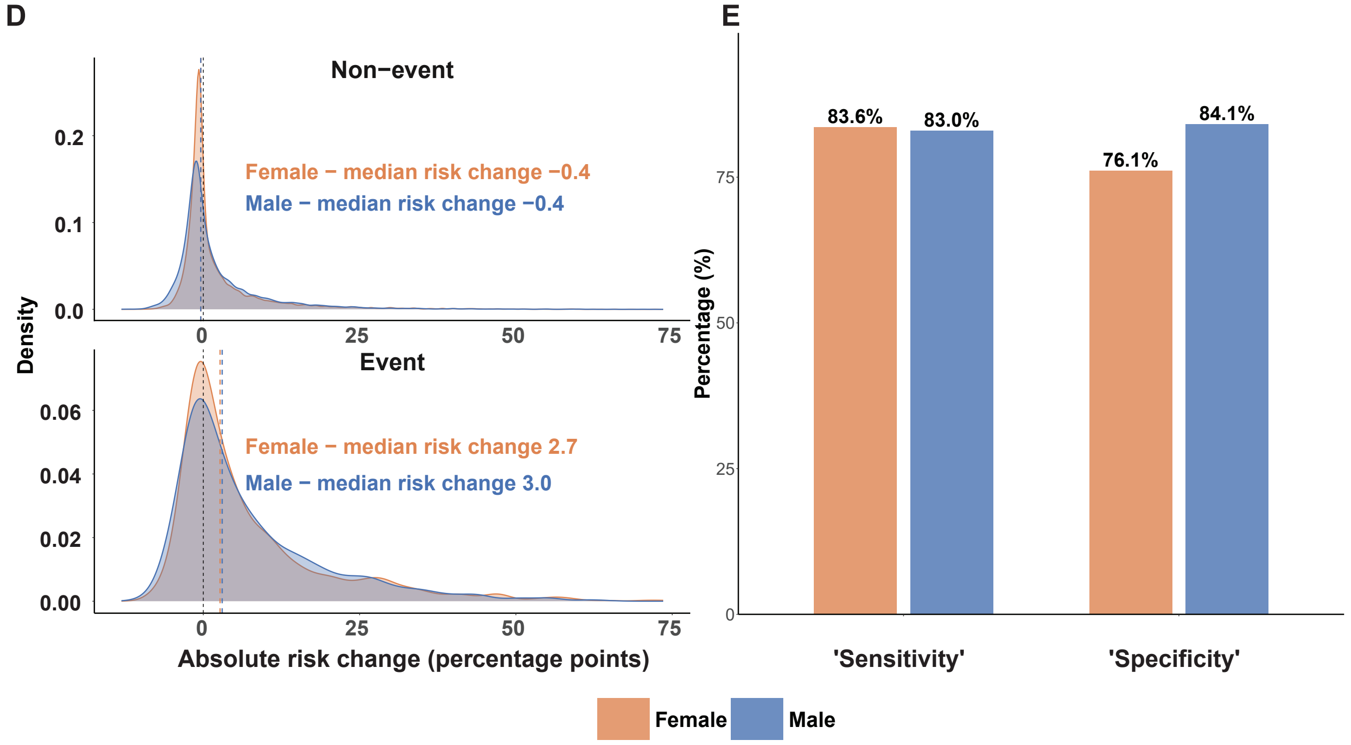
D
Non−event
Female − median risk change −0.4
Male − median risk change −0.4
Event
Female − median risk change 2.7
Male − median risk change 3.0
Density
Absolute risk change (percentage points)
0.0
0.1
0.2
0.00
0.02
0.04
0.06
0
25
50
75
E
Percentage (%)
83.6%
83.0%
76.1%
84.1%
'Sensitivity'
'Specificity'
Female
Male

**Supplementary Figure 2. Integrating IMM-AGE into PREVENT. (A)** IMM-AGE modestly improves rank discrimination beyond PREVENT. Change in c-index relative to PREVENT alone, shown separately for women, men, and the overall cohort. IMM-AGE produced consistent gains in discrimination, increasing the overall c-index by 0.0197, while the combined model achieved a similar increase of 0.0209. **(B)** Reclassification patterns following addition of IMM-AGE to PREVENT risk. Cumulative distribution of reclassified individuals by distance from the original risk threshold, stratified by event status and direction of reclassification. **(C-D)** IMM-AGE-driven risk shifts are concentrated in clinically actionable PREVENT strata. Distribution of absolute risk change after IMM-AGE incorporation, stratified by baseline PREVENT risk category, sex, and event status. Among participants who later developed cardiovascular events, upward shifts were most pronounced in intermediate- and high-risk strata, with median increases of 3.45 and 12.1 percentage points in women and 1.09 and 7.31 percentage points in men, respectively (C). Among non-events, IMM-AGE mostly shifted low- and borderline-risk individuals downward, while a subset of high-risk non-events shifted upward, consistent with elevated immune-risk profiles despite no observed event during follow-up (D). **(E)** IMM-AGE shifts individual risk in the expected direction. Distribution of per-person risk change after IMM-AGE incorporation, stratified by sex and event status. Future event cases showed median risk increases of 2.7 and 3.0 percentage points in women and men, respectively, whereas non-events showed median decreases of 0.4 percentage points in both sexes. **(F)** IMM-AGE preserves treatment-aligned reclassification performance. Sensitivity and specificity of PREVENT reclassification after IMM-AGE incorporation, defined by whether risk avoided decreasing among future event cases or increasing among non-events. IMM-AGE maintained high performance across sexes.
