## Supplementary Figure 1 for "Immune Aging is an Independent Risk Factor for Cardiovascular Disease" (from another version of this preprint: version 1, 2026-05-20)

**Supplementary Figure 1. Development and internal validation of a transcriptomic gene-ratio-based IMM-AGE signature.**

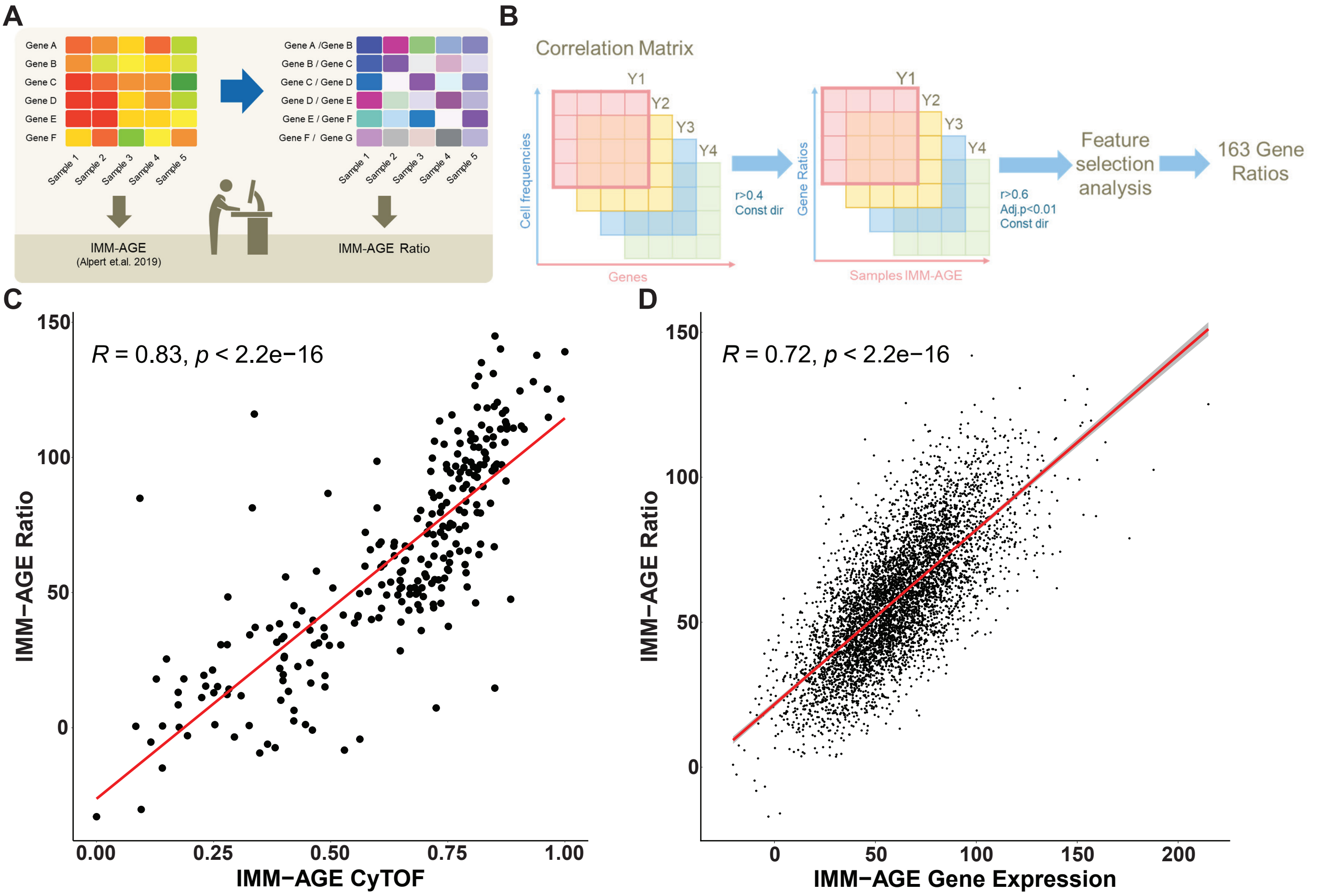

**Supplementary Figure 1. Development and internal validation of a transcriptomic gene-ratio-based IMM-AGE signature. (A)** Generalization of IMM-AGE for large-scale cohorts. A schematic outlining the development of a gene-ratio-based transcriptomic signature.
**(B)** Schematic of the gene-ratio selection workflow in the SELA discovery cohort. After filtering genes for consistent age-related signal and correlation with cytometry-defined IMM-AGE features across four years, candidate ratios were prioritized by combined significance, correlation strength and lasso-based stability selection, yielding a final signature of 163 gene ratios from 105 genes. **(C)** Correlation between CyTOF-derived IMM-AGE and gene-ratio-based estimates in SELA (n = 262; Spearman  = 0.83, P < 2.2×10-16). (D) Correlation between IMM-AGE scores derived from the original 56-gene signature and the revised 163-ratio signature in the Framingham Heart Study (n = 2,418; Spearman  = 0.72, P < 2.2×10-16).
